# Motor processivity and speed determine structure and dynamics of microtubule-motor assemblies

**DOI:** 10.1101/2021.10.22.465381

**Authors:** Rachel A. Banks, Vahe Galstyan, Heun Jin Lee, Soichi Hirokawa, Athena Ierokomos, Tyler D. Ross, Zev Bryant, Matt Thomson, Rob Phillips

## Abstract

2

Active matter systems can generate highly ordered structures, avoiding equilibrium through the consumption of energy by individual constituents. How the microscopic parameters that characterize the active agents are translated to the observed mesoscopic properties of the assembly has remained an open question. These active systems are prevalent in living matter; for example, in cells, the cytoskeleton is organized into structures such as the mitotic spindle through the coordinated activity of many motor proteins walking along microtubules. Here, we investigate how the microscopic motor-microtubule interactions affect the coherent structures formed in a reconstituted motor-microtubule system. We explore key parameters experimentally and theoretically, using a variety of motors with different speeds, processivities, and directionalities. We demonstrate that aster size depends on the motor used to create the aster, and develop a model for the distribution of motors and microtubules in steady-state asters that depends on parameters related to motor speed and processivity. Further, we show that network contraction rates scale linearly with the single-motor speed in quasi one-dimensional contraction experiments. Finally, we demonstrate that competition between motors of opposite polarity can be tuned to create various structures. In all, this theoretical and experimental work helps elucidate how microscopic motor properties are translated to the much larger scale of collective motor-microtubule assemblies.

**Significance:** In living matter, the consumption of energy at the molecular scale often results in the emergence of ordered structures that are hundreds to thousands of times greater in size. A canonical system for studying the emergence of ordered structures is the cellular cytokeleton, which primarily consists of filaments and motor proteins. Here, we use a recently developed optogenetically controlled motor-microtubule system to explore the relation between the properties of the motors that interact with microtubules, each with its own average velocity and processivity, and the structures that emerge when those motors act collectively to form asters. We find that aster size, distribution of motors within the aster, and inter-aster contraction rates are connected to these motor properties.

## 3 Introduction

A signature feature of living organisms is their ability to create beautiful, complex patterns of activity, as exemplified in settings as diverse as the famed flocks of starlings in Rome or the symmetrical and dazzling microtubule arrays that separate chromosomes in dividing cells [1]. While such organization in nature has long captured the attention of artists and scientists alike, many questions remain about how the patterns and structures created by living organisms arise. In active systems such as bird flocks or microtubule-motor arrays, energy is consumed at the local level of the individual actors, and the coordinated action of many individuals creates the much larger observed patterns. How the specific microscopic activity of each individual leads to the final large-scale assembly formed remains an open question.

The motor-microtubule system is an excellent system in which to test this question, as many motor proteins with a variety of properties, such as speeds, stall and detachment forces, processivities, and directionalities exist in nature. These motors play a variety of roles in cells; some transport cargo while others localize to distinct regions of the mitotic spindle [2–6]. Studies have investigated how the microscopic properties of these motors makes them uniquely suited to their cellular role. For example, kinesin-1’s high speed and processivity make it excellent at transporting cargo [7, 8]. However, in *in vitro* systems, kinesin-1 tetramers are able to form asters, extensile networks, and contractile networks [9–12]. Ncd (Kinesin-14) and Kif11 (Kinesin-5) have similarly been shown to form asters in *vitro*, yet it remains unclear how the properties of these motors affect the structure and dynamics of the assemblies created [9, 13].

In this work, we create motor-microtubule structures with a variety of motors and develop theoretical models to connect the motor properties to the organization and dynamics of the assemblies. Our recently developed optogenetic *in vitro* motor-microtubule system demonstrated the formation of asters and other contractile networks with kinesin-1 (K401) upon light activation [12]. We now show how this system can be made to work with kinesin-5 (Kif11) and kinesin-14 (Ncd), and form asters of varying sizes with each motor. Further, we develop a theoretical model connecting the distribution of motors and microtubules in asters; calculated distributions depend on the motor properties and fit with our experimental data. By using motors with different speeds, we find that contraction rates in quasi one-dimensional microtubule networks directly depend on the single-motor velocity. Finally, we set up competition between motors of opposite polarity, and find that the final structure formed depends on the properties and concentrations of each motor. This theoretical and experimental work sheds light on the ways that microscopic motor properties are reflected in the thousand-fold larger length scale of motor-microtubule assemblies.

## 4 Results

### 4.1 Aster Size Depends on Motor Used

We build on the foundational work that demonstrated the ability to control motor-microtubule systems with light [12] to consider a new set of motors with different fundamental properties. In brief, kinesin motors are fused to the light-dimerizable pair iLid and micro. In the absence of light, motor dimers walk along microtubules but do not organize them; upon activation with light, the motor dimers couple together to form tetramers, crosslinking the microtubules they are walking along as shown in Fig. 1(A). Projecting a disk pattern of light on the sample results in the formation of an aster.

**Figure 1:**
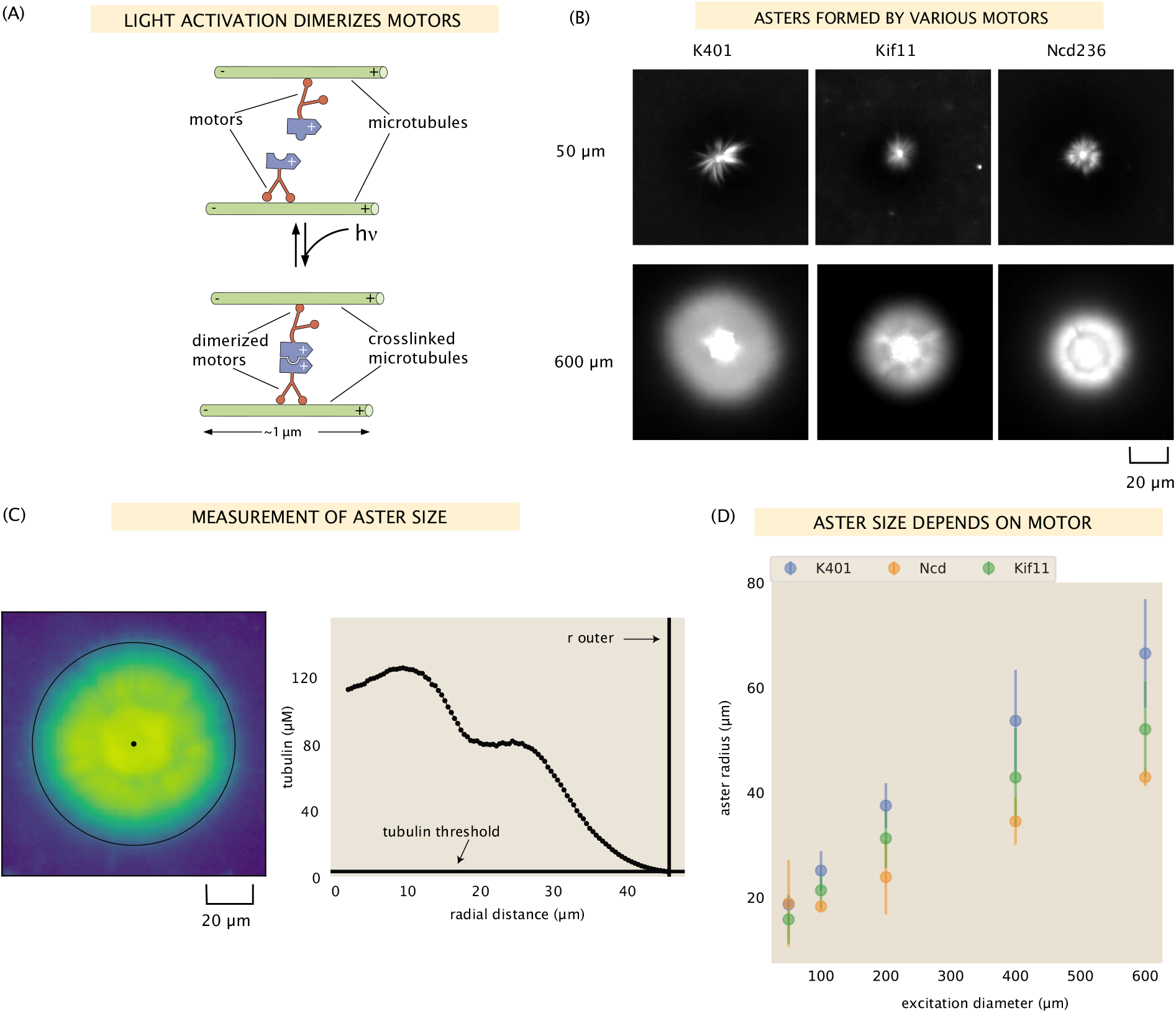
Aster size depends on motor used. (A) Motor heads are fused to optogenetic proteins such that activation with light causes the formation of motor tetramers (dimer of dimers). Motors are shown walking towards the microtubule plus-end. K401 and Kif11 walk in this direction, however Ncd is minus-end directed. (B) Images of the microtubule fluorescence for asters formed with each of the motors excited with a disk either 50 μm or 600 μm in diameter. (C) Image of the microtubule fluorescence from an aster with the measured size represented with the outer black circle. The plot on the right shows the radial microtubule concentration as revealed by fluorescence intensity; the threshold concentration used to determine the aster size is shown as a black horizontal line. (D) Mean aster size (n ≈ 5 asters for each condition) for the three motors and different excitation diameters; the error bars represent the standard deviation.

In this work, we aim to determine how the properties of the motor affect the resulting structures. While experiments with this system were previously performed with *D. melanogaster* kinesin-1 motors (K401) [12], in the present work, we investigate if other kinesin motor species with different intrinsic properties such as speed and processivity would lead to light-inducded microtubule organization. Towards this end, we use the same light-dimerizable scheme to form microtubule structures with two other motors: Ncd (D. *melanogaster* kinesin-14) and Kif11 *(H. sapiens* kinesin-5). The single-molecule properties of all three motors we use are summarized in Table 1. We measure the speed of each motor species by gliding assays (SI section 9.9); the processivities are based on literature values. Further, we fluorescently label the motors using mVenus or mCherry to visualize the motors and microtubules in separate imaging channels within the same assay (SI section 1).

**Table 1:** Motor proteins used and their properties

| Motor | Speed | Processivity | Direction |
| --- | --- | --- | --- |
| K401 | $\approx 600$ nm/s | $\approx 100$ steps [14] | plus [8] |
| Ncd236 | $\approx 115$ nm/s | not processive [15–17] | minus [18] |
| Kif11(513) | $\approx 70$ nm/s | $\approx 10$ steps [19] | plus [19] |

As seen in Fig. 1(B) and Fig. S4, each of these motors is able to form asters of varying sizes in our system. It was previously unclear whether there were limits to speed, processivity, or stall force that might prevent any of these motors from forming asters in our light-controlled system, although Ncd has previously been shown to form asters as constitutive oligomers [20, 21]. We found that all were able to form asters upon illumination by various excitation diameters ranging from 50 to 600 μm. In order to measure the size of the asters, we used the distribution of fluorescently labeled microtubules, which peaks in the center of the aster and generally decreases moving outward. We defined the outer radius of the aster as the radius at which the microtubule fluorescence is twice the background microtubule concentration (see Fig. 1(C) for an example aster outer radius determination). We found this method to agree well with a visual inspection of the asters (Fig. S4).

We find that aster size increases with excitation diameter (Fig. 1(B), (D)), consistent with what was shown by Ross et al. for K401 [12]. Interestingly, we find that the size of the asters also depends on the motor used. For each excitation diameter, except for the 50 μm case, K401 formed the largest asters and Ncd formed the smallest, with Kif11 producing asters of intermediate size (Fig. 1(D)). What is it about the different motors that confers these different structural outcomes? We found that this trend correlates with motor processivity; K401 is the most processive, followed by Kif11, and then Ncd. This is similar to the findings in [9], in which ‘intensity’ of aster formation was related to motor processivity. Other factors could also be contributing to aster size such as the ratio of microtubules to motors as was suggested by [9] but not investigated in the present work, or the motor stall force.

### 4.2 Spatial Distribution of Motors in Asters

The nonuniform distribution of filaments and motors in an aster is a key feature of its organization and has been the subject of previous studies. In these studies, continuum models were developed for motor-filament mixtures which predicted the radial profile of motors in confined two-dimensional systems [22–25]. A notable example is the power-law decay prediction by Nédélec et al., who obtained it for a prescribed organization of microtubules obeying a 1/*r* decay law [22]. Measuring the motor profiles in asters formed in a quasi-two-dimensional geometry (with the *z* dimension of the sample being only a few microns deep) and fitting them to a power-law decay, the authors found a reasonable yet noisy match between the predicted and measured trends in the decay exponent.

In our work, we also develop and test a minimal model that predicts the motor profile from the microtubule distribution and the microscopic properties of the motor. In contrast to the earlier study [22], asters formed in our experiments are three-dimensional due to the much larger depth of the flow cells (roughly 100 μm). While the largest asters are likely partially compressed in the *z*-direction, we assume that this effect does not significantly alter the protein distributions in the central *z*-slice and hence, for modeling purposes we consider our asters to be radially symmetric, as depicted schematically in Fig. 2(A). Our modeling applies to locations outside the central disordered region (called aster ‘core’) with a typical radius of ≈ 15 μm, beyond which microtubules have a predominantly polar organization (see SI section 9.10 for the discussion of the two aster regions and an example PolScope image that demonstrates their distinction).

**Figure 2:**
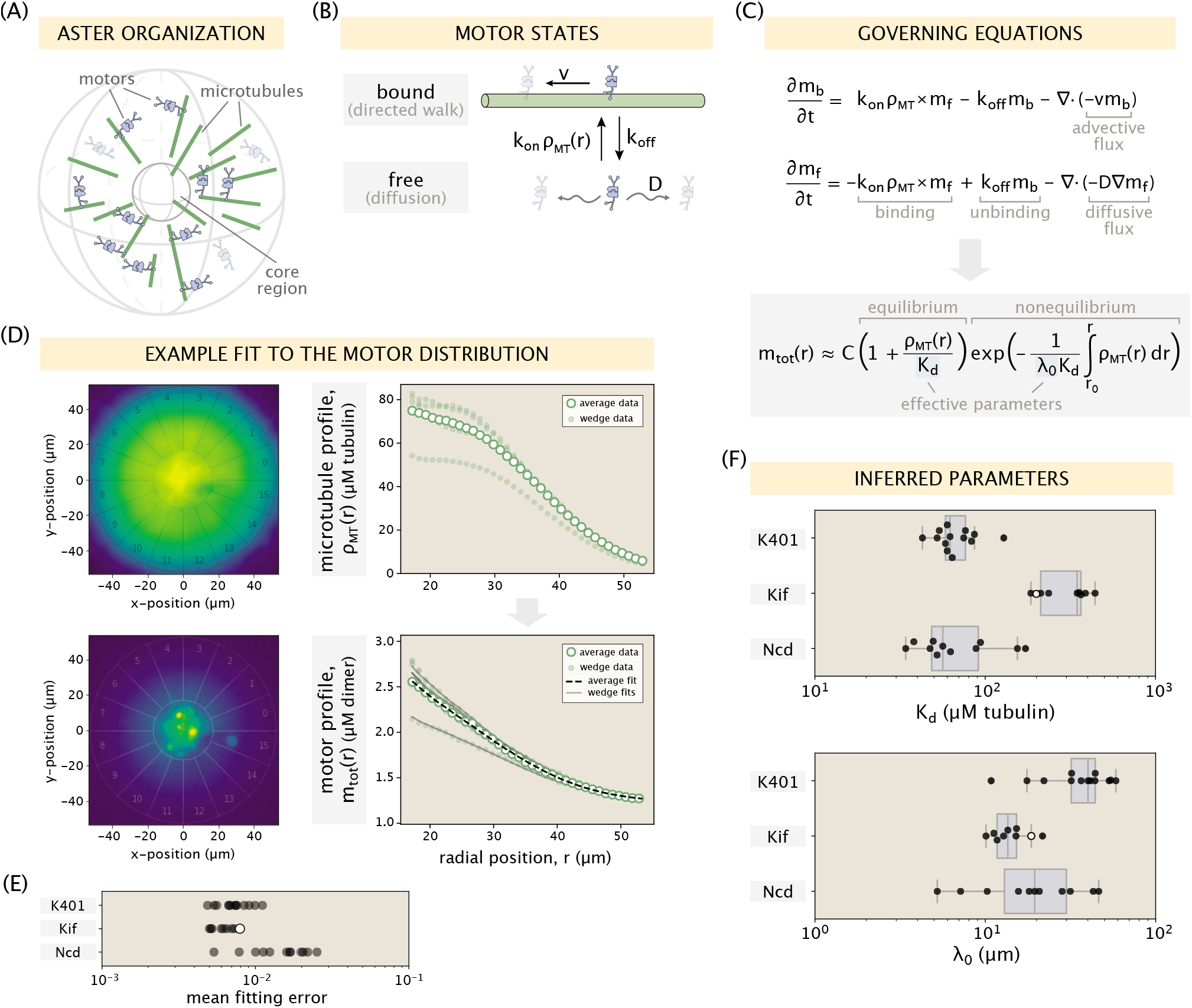
Modeling the motor distribution. (A) Schematic of the radial microtubule organization in an aster. Modeling applies to locations outside the disordered core region at the aster center. Components of the schematic are not drawn to scale. (B) Motor states and transitions between them. (C) Governing equations for the bound and free motor populations, along with our solution for the total motor distribution at steady state, expressed via effective parameters *K*_d_ = *k*_off_/*k*_on_ and λ_0_ = *D*/*v* (see SI section 9.15.1 for details). (D) Demonstration of the model fitting procedure on an example Kif11 aster. Fits to the average motor profile as well as to 5 out of 16 wedge profiles are shown. The outlier case with a lower concentration corresponds to wedge 13 in the fluorescence images. (E) Mean fitting errors for all asters calculated from the fits to the wedge profiles. The error is defined as the ratio of the mean residual to the concentration value at the inner boundary. (F) Inferred parameters *K*_d_ and λ_0_ grouped by the kind of motor. Box plots indicate the quartiles of the inferred parameter sets. The fitting error and the inferred parameters for the Kif11 aster in panel (D) are shown as white dots in panels (E) and (F).

Similar to the treatment in earlier works [22, 24, 25], we introduce two states of the motor – an unbound state where the motor can freely diffuse with a diffusion constant *D* and a bound state where the motor walks towards the aster center with a speed *v* (Fig. 2(B)). In the steady-state of the system, which we assume our asters have reached at the end of the experiment, microtubules on average have no radial movement and hence, do not contribute to motor speed. We denote the rates of motor binding and unbinding by *k*_on_ and *k*_off_, respectively. When defining the first-order rate of motor binding, namely, *k*_on_*ρ*_MT_(*r*), we explicitly account for the local microtubule concentration *ρ*_MT_(*r*) extracted from fluorescence images. This is unlike the previous models which imposed specific functional forms on the microtubule distribution (e.g., a constant value [23, 24], or a power-law decay [22]), rendering them unable to capture the specific features often seen in our measured microtubule profiles, such as the presence of an inflection point (see Fig. 1(C) for an example).

The governing equations for the bound (*m*_b_) and free (*m*_f_) motor concentrations are shown in Fig. 2(C). They involve binding and unbinding terms, as well as a separate flux divergence term for each population. Solving them at steady state, we arrive at an equation for the total local concentration of motors defined as *m*_tot_(*r*) = *m*_b_(*r*) + *m*_f_(*r*). The derivation of this result can be found in SI section 9.15.1. As seen in the equation for *m*_tot_(*r*) (Fig. 2(C)), knowing the microtubule distribution *ρ*_MT_(*r*) along with two effective microscopic parameters, namely, the effective dissociation constant *K*_d_ = *k*_off_/*k*_on_ and the length scale λ_0_ = *D*/*v*, one can obtain the motor distribution up to a multiplicative constant (*C* in the equation). Note that in the special case where the motors do not move (*v* → 0 or λ_0_ → ∞), the exponential term becomes 1 and an equilibrium relation between the motor and microtubule distributions dependent only on *K*_d_ is recovered.

To test this model, we extract the average radial distributions of microtubule and motor concentrations for each aster. Then, using the microtubule profile as an input, we fit our model to the motor data and infer the effective parameters *K*_d_ and λ_0_ (see SI sections 9.15.2 and 9.15.3 for details). A demonstration of this procedure on an example Kif11 aster is shown in Fig. 2(D) where a good fit to the average motor data can be observed. As a validation of our inference method, we additionally extract the radial concentration profiles inside separate wedges of the aster and show that they can be accurately captured by only choosing an appropriate multiplicative constant *C* for each wedge, while keeping the pair (*K*_d_, λ_0_) inferred from average profile fixed (fits to 5 out of 16 different wedge profiles are shown in Fig. 2(D) for clarity). The fitting error for other asters is similarly low (Fig. 2(E), see Fig. S10 for the collection of fitted profiles).

Plotting the inferred parameters *K*_d_ and λ_0_ from all fits (Fig. 2(F)), we find that they are clustered around single values for each motor type and vary between the motors. Based on the single-molecule motor properties in Table 1 and the reported motor binding rates [26], our expectation was that the *K*_d_ values for Kif11 and K401 would have a ratio of ≈ 4.6: 1 (see SI section 9.15.4), while *K*_d_ for Ncd would be the highest due to its non-processivity. The ratio of median inferred *K*_d_ values for Kif11 and K401 is ≈ 5.6: 1 – close to our expectation. However, the inferred *K*_d_ values for Ncd are low and comparable to those for K401. One possible resolution of this discrepancy comes from the finding of an *in vitro* study suggesting a substantial increase in the processivity of Ncd motors that act collectively [27]. Specifically, a pair of Ncd motors coupled through a DNA scaffold was shown to have a processivity reaching 1 μm (or, ≈ 100 steps) – a value close to that reported for K401 motors. A highly processive movement was similarly observed for clusters of HSET (human kinesin-14) [28] and plant kinesin-14 motors [29]. This collective effect, likely realized for Ncd tetramers clustered on microtubules in highly concentrated aster structures, is therefore a possible cause for the low inferred values of their effective *K*_d_. We also note that while a similar collective effect on processivity was observed for K401 motors [27], it is far less dramatic due to their already large single-motor processivity, and therefore would have a small effect on the effective *K*_d_.

Next, looking at the inference results for the λ_0_ parameter (Fig. 2(F)), we can see that Kif11 and Ncd motors have an average λ_0_ value of ≈ 10 – 20 μm, while the average value for K401 motors is ≈ 40 μm. From the measured diffusion coefficient of *D* ≈ 1 μm^2^/s for tagged kinesin motors [7] and the single-molecule motor speeds reported in Table 1, our rough estimate for the λ_0_ parameter for Kif11 and Ncd motors was ≈ 10 — 15 μm, and ≈ 2 μm for K401. While the inferred values for the two slower motors are well within the order-of-magnitude of our guess, the inferred λ_0_ for the faster K401 motor is much higher than what we anticipated. This suggests a significant reduction in the effective speed. One contributor to this reduction is the stalling of motors upon reaching the microtubule ends. Recall that in our model formulation (Fig. 2(B)) we assumed an unobstructed walk for bound motors. Since the median length of microtubules (≈ 1.6 μm) is comparable to the processivity of K401 motors (≈ 1 μm), stalling events at microtubule ends will be common, leading to a reduction of their effective speed in the bound state by a factor of ≈ 1.5 (see SI section 9.15.5 for details). This correction alone, however, is not sufficient to capture the factor of ≈ 25 discrepancy between our inference and the estimate of λ_0_. We hypothesize that an additional contribution may come from the jamming of K401 motors in dense aster regions. This is motivated by the experiments which showed that K401 motors would pause when encountering obstructions during their walk [30, 31]. In contrast, for motors like Ncd and Kif11 which take fewer steps before unbinding and have a larger effective *K*_d_, jamming would have a lesser effect on their effective speed as they would unbind more readily upon encountering an obstacle. Overall, our study shows that the minimal model of motor distributions proposed in Fig. 2 is able to capture the distinctions in aster structure through motor-specific effective parameters, although more work needs to be done to explain the emergence of higher-order effects such as motor clustering and jamming, and their contribution to these effective parameters.

Our model also provides insights on the observation that the distribution of microtubules is generally broader than that of the motors. This feature can be observed by comparing the two example profiles in Fig. 2(D), and it also holds for the profiles extracted from other asters (Fig. S10). In SI section 9.15.6, we use our model to demonstrate this feature in a special analytically tractable case, and discuss its generality across asters in greater detail. We found that the relative width of the motor distribution compared to the microtubule distribution is fairly constant among asters, with their difference being the largest for Ncd motors, consistent with our model predictions. This relationship between the shapes of the distributions may be an important factor in the spatial organization of end-directed motors in the spindle where their localization to the spindle pole is of physiological importance.

### 4.3 Contraction Rate Scales with Motor Speed

Besides the steady-state structure of motor-microtubule assemblies, it is also of great interest to understand their dynamics. Ross et al. demonstrated the formation of quasi one-dimensional contractile networks by creating two asters that are initially separated by a given distance, then activating a thin rectangular region between them to form a network connecting the asters that pulls them together [12]. Example images of microtubule fluorescence during one of these experiments are shown in Fig. 3(A). In these experiments, an increase in maximum aster merger speed with greater initial separation was observed. We aimed to confirm this behavior with our various motors and to test the relationship between aster merger speed and single motor speed.

**Figure 3:**
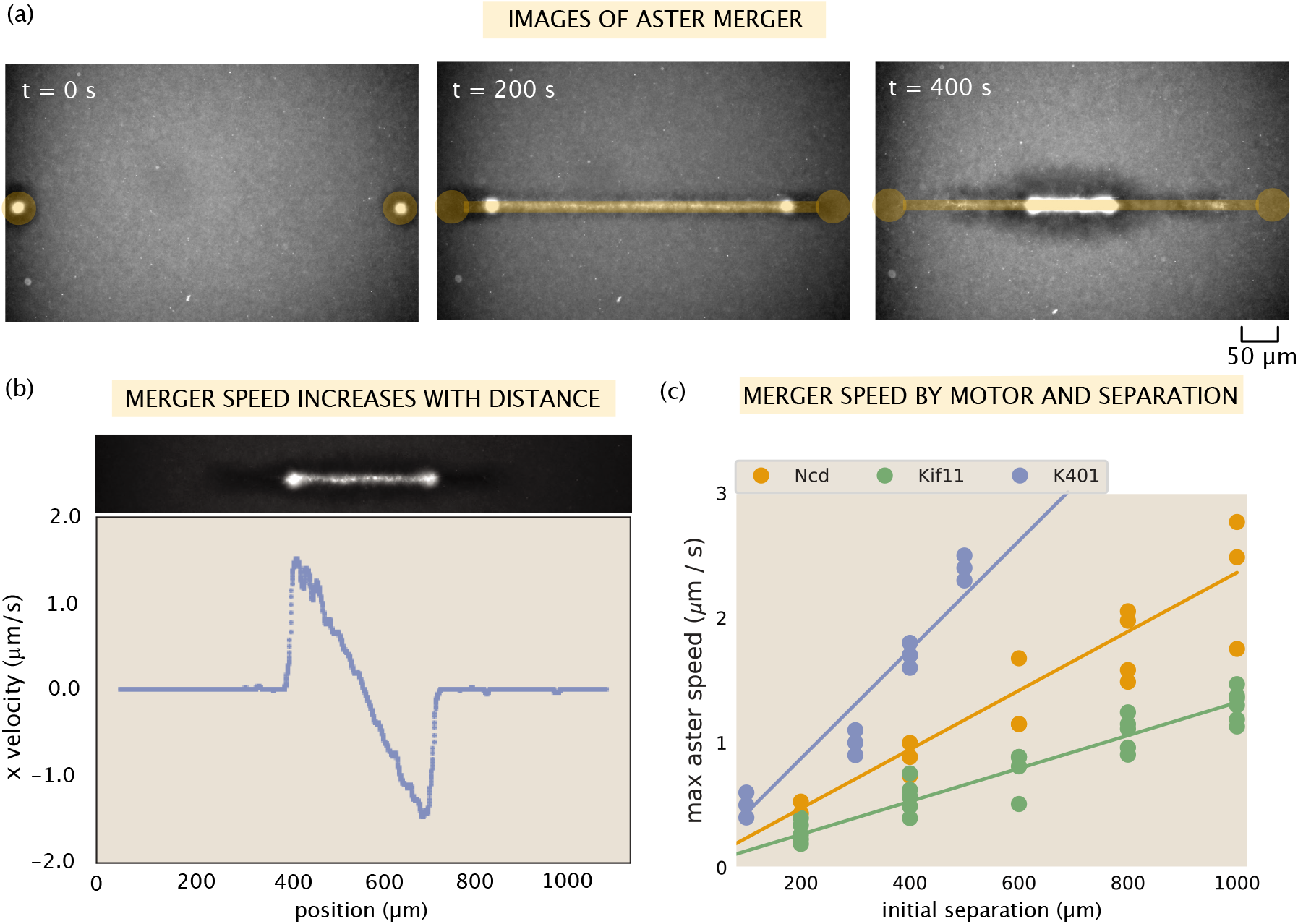
Contractile speeds in motor-microtubule networks scale with network size and motor speed. (A) Images of microtubule flourescence during aster merger. Regions of light activation are shown in orange. (B) Example profile of speeds in an aster merger as a function of linear distance. Each dot is the mean speed measured at that x-position within the network. (C) Maximum merger speed, measured at the ends of the network for each initial separation and motor. Each dot is a single experiment and the lines are best fits to the data.

First, we tested the relationship between distance and speed in our experiments. Using optical flow to measure the contraction speed throughout the network, we observe a linear increase in contractile speed with distance from the center of the network, as shown in Fig. 3(B). This relationship suggests that the contractile network can be thought of as a series of connected contractile units. Independent contraction of each unit would generate the observed linear increase in speed because more contractile units are added with distance from the center of the network.

Next, we investigated how contractile speeds vary by motor in aster merger experiments. We repeated aster merger experiments with each motor and with various initial separations between the asters. Fig. 3(C) shows the results, where each point represents the maximum aster speed measured in a single experiment by tracking the aster, and the lines are linear fits to the data for each motor (see SI section 9.12 for details on measuring the aster speeds). Interestingly, the ratios of the slopes of these lines match the ratios of motor speeds from Table 1. For example, the slope of the best fit line for Ncd is ≈ 0.0023 s^-1^ and the best fit slope for Kif11 is ≈ 0.0013 s^-1^. The ratio of these (Ncd/Kif11) is ≈ 1.8, which closely matches the ratio of their single motor speeds (≈ 1.6). Similar calculations can be done with these two motors and K401, with the same result. Thus, we conclude that the rate of contraction in the network is set by the motor speed and the increase of network speed with distance is due to adding more connected contractile units.

### 4.4 Motor Properties Determine Outcomes of Motor Competitions

After examining how the properties of individual species of motors affect the structures they generate, we asked what would happen when two motors of opposing polarities are active in the same system. This is inspired by the situation in cells, in which many motors are present and walking towards different poles of the microtubules. Previously, ‘tug-of-war’ experiments have been performed in which specified numbers of motors are attached to microtubules and the microtubule movement is measured [32]. These experiments showed that collections of weaker motors working together can overcome stronger, faster motors [33]. In addition, experiments were performed that tested for spontaneous aster formation in the presence of both Eg5 and XCTK2 (*Xenopus laevis* kinesin-5 and −14). They found that XCTK2 always ‘won’ and formed asters, while with an engineered motor where the Eg5 motor domain was replaced with that of kinesin-1, XCTK2 and kinesin-1 sorted to form a mixture of asters with plus-end centers and minus-end centers [21]. In addition, Nédélec simulated heteromotor complexes to explore the range of steady states that could be achieved when these motors acted on already formed asters, and found that the result depended on the relative velocities of the two motors [34]. Our system offers a unique opportunity to test the effects of motor competition in aster formation by directly dimerizing opposing motors to each other.

We first tested if an aster could be formed with heterodimers of K401 and Ncd (the second motor pair in Fig. 4(A)). Previous experiments were performed with competing K401-K401 and Ncd-Ncd dimers, and resulted in alternating plus and minus-end centered asters [35]. In our system, we form K401-Ncd heterodimers, which results in bundles of microtubules, but we did not observe overall contraction and reorganization to form an aster (Movie S1). Images of the microtubule and YFP- Ncd fluorescence are shown in the middle row of Fig. 4(B). Thus, neither motor dimer was able to ‘win’ to form an aster in this case.

**Figure 4:**
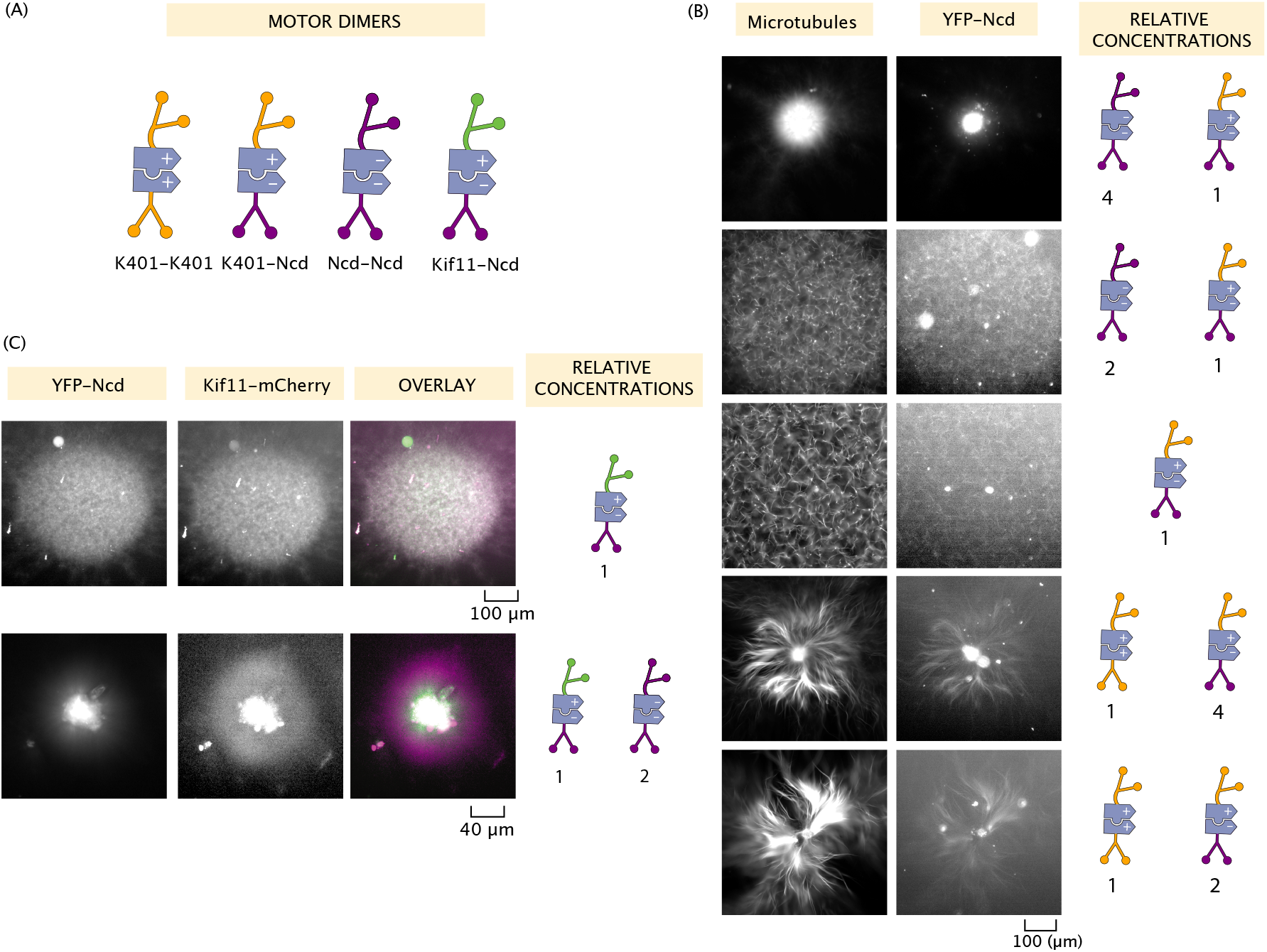
Motor competition modulates structure formation. (A) Cartoon of the various motor dimers present. Homodimers as in the previous experiments (K401-K401 and Ncd-Ncd) and heterodimers of K401-Ncd and Kif11-Ncd are formed. (B) Various combinations of K401 and Ncd motors and the resultant structures. Left – microtubule fluorescence images. Right – Ncd fluorescence images. (C) Asters formed with combinations of Ncd and Kif11 motors. In the overlay, Ncd fluorescence is in green, Kif11 is in magenta. For (B) and (C), results are shown for excitation with a 600 μm diameter disk.

Next, we tested if we could make an aster form by adding extra K401-K401 or Ncd-Ncd dimers. The results of these experiments are shown in Fig. 4(B), with the top having the most Ncd relative to K401 and moving downward, this ratio decreases. Upon the addition of even a small amount of K401-K401 dimers, (1:4 K401-K401 to K401-Ncd), K401 was able to form an aster (Movie S2). The addition of even more K401-K401 dimers increased the contraction rate of formation, indicating that the opposing force exerted by Ncd was more easily overcome (Movie S3). Interestingly, a larger amount of Ncd-Ncd dimers were required to make Ncd ‘win’. At 2:1 Ncd-Ncd to K401-Ncd, the network still failed to contract, although a brief contractile phase was observed but then stalled (Movie S4). Ncd was finally able to form an aster with four times as many Ncd-Ncd dimers as heterodimers (Movie S5). This difference in the relative amount of motors required to make K401 and Ncd dominated asters correlate with the larger speed and processivity of K401 motors.

We performed similar experiments with YFP labeled Ncd and mCherry labeled Kif11, allowing us to visualize each motor. Similarly, with only the heterodimer, bundles of microtubules were formed but no overall contraction was observed (Fig. 4(C), top row, and Fig. S7(A)). With a ratio of only 2:1 Ncd-Ncd to Kif11-Ncd, however, an aster with Ncd at the center was formed. This can be seen by comparing the YFP and mCherry fluorescence in the bottom row of Fig. 4(C). While the YFP fluorescence is highly localized in the center of the aster, the mCherry fluorescence is more prevalent in the arms of the aster. The relative fluorescence intensity of each is plotted in Fig. S7(B), demonstrating that the ratio of Kif11 motors relative to Ncd motors increases moving radially outward from the center of the aster. This indicates that the minus ends of the microtubules are at the center, and the plus ends are pointing outwards which Kif11 walks towards. Comparing the ratio of motors required for Ncd to ‘win’, Ncd was able to overcome the competing forces of Kif11 at a lower concentration than it did against K401. This is likely due to the lower speed and processivity of Kif11 compared to K401.

## 5 Discussion

In this work, we examined how the properties of kinesin motors determine the mesoscopic properties of the structures they create. The way in which quantities such as motor speed and processivity govern the nature of the resulting motor-microtubule structures has been an open question. Previous attempts have been made to address this question, however most of these are in the context of a single motor without varying the speed or processivity and developing a model that fits the properties they measure [22–25]. By varying motor speeds, processivities, and directionalities, we were able to quantify and model how these microscopic parameters connect to the properties of mesoscopic structures.

We demonstrate light-controlled aster formation with three different motors. Interestingly, the final aster size from a given illumination region varied depending upon which motor was used. Our leading hypothesis is that the key control variable is the processivity of the motors. Future work needs to be done to understand this effect and build models to explain it. Early work by Surrey et al. found that processivity affected the ‘intensity’ of aster formation in simulations, which may be related to our observations, but to the best of our knowledge no model of this effect has been developed [9]. Further, we assess the distribution of motors and microtubules in the asters we form and develop a model of the steady-state aster that predicts the motor distribution given the measured microtubule distribution, with parameters that relate to the motor speed and processivity. Interestingly, the parameters we infer differ from those we would expect from the single molecule properties of the motors, indicating that higher order effects such as increased processivity of collections of motors, are playing important roles. In addition, we measure contraction speeds in pseudo-one-dimensional networks and find that the speeds are related to the single motor speed. Finally, we show how competition between motors of opposite polarity can lead to tunable structure outcomes. The question of how individual microscopic properties are translated to the hundreds to thousands of times larger scale of the assembly has been an open question in the field of active matter. Our well controlled motor-microtubule system provides an excellent arena in which to investigate this question and our work takes a step towards a mechanistic understanding of motor-microtubule assemblies.

## 6 Acknowledgements

We are grateful to Dan Needleman, Madhav Mani, Peter Foster, and Ana Duarte for fruitful discussions. We acknowledge support from the NIH through grant 1R35 GM118043-01; the John Templeton Foundation as part of the Boundaries of Life Initiative through grants 51250 and 60973; the Foundational Questions Institute and Fetzer Franklin Fund through FQXi 1816.

## 7 Data Sharing Plans

All code will be stored on our Github Repository and additional data will be housed on the Caltech Data Repository.

## 8 Competing Interests

The authors declare no competing interests.

**Table S1:**
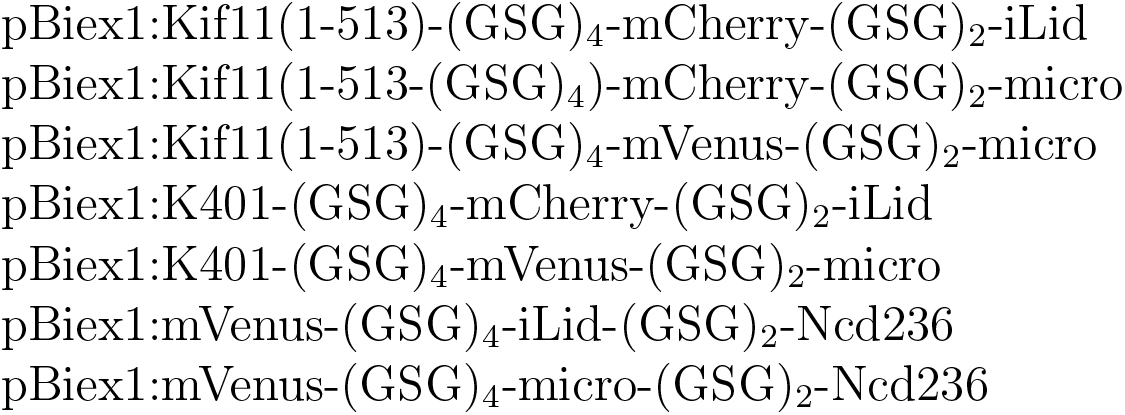
Plasmids used

## 9 Materials and Methods

### 9.1 Cloning of Motor Proteins

Human kinesin-5 (Kif11/Eg5) 5-513 was PCR amplified from mCherry-Kinesin11-N-18 plasmid (gift from Michael Davidson, Addgene # 55067). This fragment was previously shown to form functional dimers [19]. Kinesin 1 1-401 (K401) was PCR amplified from pWC2 plasmid (Addgene # 15960). Ncd 236-701 was PCR amplified from a plasmid gifted by Andrea Serra-Marques.

The optogenetic proteins, iLid and micro were PCR amplified from pQE-80L iLid(addgene # 60408, gift from Brian Kuhlman) and pQE-80L MBP-SspB Micro (addgene # 60410). mCherry was PCR amplified from mCherry-Kinesin11-N-18 and mVenus was PCR amplified from mVenus plasmid (addgene # 27793).

Constructs were assembled by Gibson assembly of the desired motor protein, optogenetic protein, and fluorophore in order to make the plasmids listed in Table S1.

### 9.2 Protein Expression and Purification

Protein expression and purification was done in SF9 cells. Cells were seeded at a density of 1,000,000 cells per mL in a 15 mL volume and transiently transfected with the desired plasmid using Escort IV transfection reagent, then incubated for 72 hours before purification. Cells were collected for purification by centrifugation at 500 g for 12 minutes, and the pellet was resuspended in lysis buffer (200 mM NaCl, 4 mM MgCl2, 0.5 mM EDTA, 1.0 mM EGTA 0.5 % Igepal, 7 % Sucrose by weight, 20 mM Imidazole pH 7.5, 10 μg/mL Aprotinin, 10 μg/mL Leupeptin, 2 mM ATP, 5 mM DTT, 1 mM PMSF) and incubated on ice for 30 minutes. The lysate was then clarified by centrifugation at 200,000 g for 30 minutes at 4°C. Clarified supernatant was incubated with 40 μL anti-FLAG M2 affinity gel (Sigma-Aldrich A2220) for 3 hours at 4°C. To wash out unbound protein, the resin (with bound protein) was collected by centrifugation at 2,000 g for 1 minute, the supernatant was removed and the resin was washed with wash buffer (for Ncd and Kif11: 150 mM KCl, 5 mM MgCl2, 1 mM EDTA, 1 mM EGTA, 20 mM Imidazole pH 7.5, 10 μg/mL Aprotinin, 10 μg/mL Leupeptin, 3 mM DTT, 3mM ATP; for K401: M2B with 10 μg/mL Aprotinin, 10 μg/mL Leupeptin, 3 mM DTT, 3mM ATP). This was repeated two more times with decreasing ATP concentration (0.3mM and 0.03mM ATP) for a total of three washes. After the third wash, about 100 μL supernatant was left and the bound protein was eluted by incubation with 10 μL FLAG peptide (Sigma-Aldrich F3290) at 4°C for 3 hours. The resin was then spun down by centrifugation at 2,000 g for 1 minute and the supernatant containing the purified protein was collected. Purified protein was then concentrated to a volume of 10-20 μL by centrifugation in mini spin filters (Millipore 50 Kda molecular weight cut-off). Protein was kept at 4°C and used the same day as purification or stored in 50% glycerol at −20°C for longer storage. Protein concentration was determined with QuBit Protein Assay Kit (Thermo Fisher Q33212).

### 9.3 Microtubule Polymerization

Microtubules were polymerized as reported previously ([12] and originally from the Mitchison lab website [36]). In brief, 75 μM unlabeled tubulin (Cytoskeleton) and 5 μM tubulin-AlexaFluor647 (Cytoskeleton) were combined with 1mM DTT and 0.6mM GMP-CPP in M2B buffer and incubated spun at 300,000 g to remove aggregates, then the supernatant was incubated at 37°C for 1 hour to form GMP-CPP stabilized microtubules.

### 9.4 Microtubule Length

The GMP-CPP stabilized microtubules were imaged with TIRF microscopy to determine their length. A flow chamber was made using a KOH cleaned slide, KOH cleaned coverslip (optionally coated with polyacrylamide) and parafilm cut into chambers. The flow cell was incubated with poly-L-lysine for 10 minutes, washed with M2B, then microtubules were flown in. The chamber was sealed with Picodent and imaged with TIRF microscopy.

Microtubules were segmented using home-written Python code and histogrammed to determine the distribution of microtubule lengths (Fig. S1).

**Figure S1:**
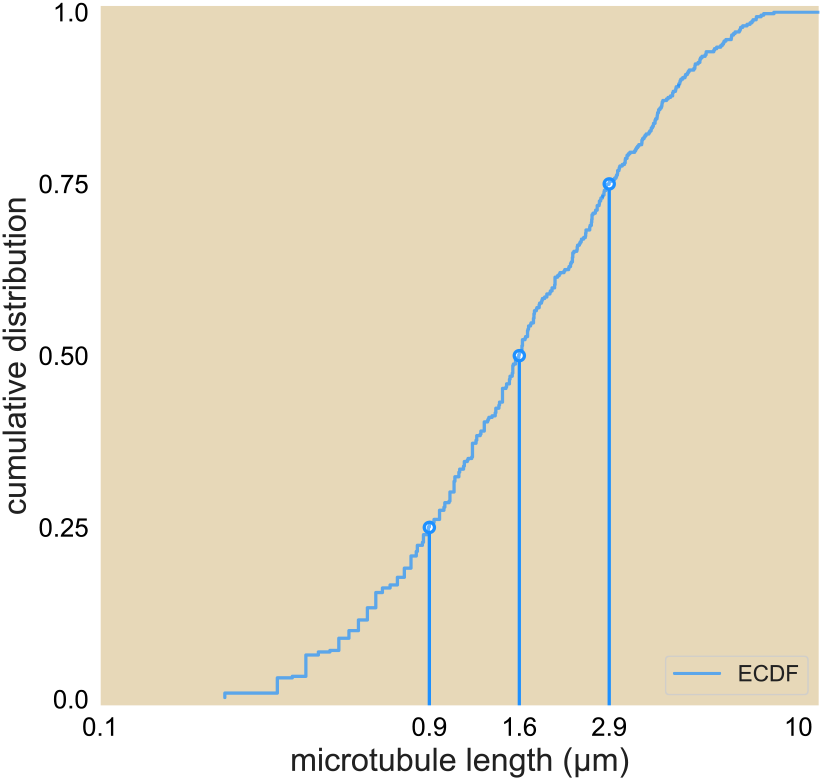
Cumulative distribution of microtubule lengths. The 25, 50, and 75% length are marked.

### 9.5 Sample Chamber Preparation

Slides and coverslips were cleaned with Helmanex, ethanol, and KOH, silanized, and coated with polyacrylamide as in [12]. Just before use, slides and coverslips were rinsed with MilliQ water and dried with compressed air. Flow chambers (3 mm wide) were cut out of Parafilm M and melted using a hotplate at 65 °C to seal the slide and coverglass together, forming chambers that are ≈ 70 — 100 μm in height and contain ≈ 7 μL.

### 9.6 Reaction Mixture Preparation

The reaction mixture consisted of kinesin motors (~ 250nM), microtubules (~ 1 μM tubulin), and energy mix that contained ATP, an ATP recycling system, a system to reduce photobleaching, F-127 pluronic to reduce interactions with the glass surfaces, and glycerol [12]. To prevent pre-activation of the optogenetic proteins and photobleaching of the fluorophores, the motors and microtubules were always handled in a dark room where wavelengths of light below 520 nm were blocked with a filter or a red light was used to illuminate. The reaction mixture was prepared right before loading into the flow cell and then sealed with Picodent Speed.

### 9.7 Microscope Instrumentation

We performed the experiments with an automated widefield epifluorescence microscope (Nikon TE2000). We custom modified the scope to provide two additional modes of imaging: epi-illuminated pattern projection and LED gated transmitted light. We imaged light patterns from a programmable DLP chip (EKB TEchnologies DLP LightCrafterTM E4500 MKIITM Fiber Couple) onto the sample through a user-modified epi-illumination attachment (Nikon T-FL). The DLP chip was illuminated by a fiber coupled 470 nm LED (ThorLabs M470L3). The epi-illumination attachment had two light-path entry ports, one for the projected pattern light path and the other for a standard widefield epi-fluorescence light path. The two light paths were overlapped with a dichroic mirror (Semrock BLP01-488R-25). The magnification of the epi-illuminating system was designed so that the imaging sensor of the camera (FliR BFLY-U3-23S6M-C) was fully illuminated when the entire DLP chip was on. Experiments were run with Micro-Manager [37], running custom scripts to controlled pattern projection and stage movement.

### 9.8 Activation and Imaging Protocol

For the experiments in which we make asters with excitation disks of different sizes, we use five positions within the same flow cell simultaneously in order to control for variation within flow cells and over time. Each position is illuminated with a different sized excitation region: 50, 100, 200, 400, or 600 μm diameter cylinder. Each position was illuminated with the activation light for ~ 50 — 200 ms and both the microtubules (Cy5 labelled) and motors (mVenus labelled) were imaged at 10X magnification every 15 seconds. After an hour of activation, a z-stack of the microtubule and motor fluorescence throughout the depth of the flow chamber was taken at 5 μm increments in each position. Typically, one experiment was run per flow chamber. We placed the time limitations on the sample viewing to minimize effects related to cumulative photobleaching, ATP depletion, and global activity of the light-dimerizable proteins. After several hours, inactivated “dark” regions of the sample begin to show bundling of microtubules.

For the aster merger experiments, two 50 μm disks are illuminated at different distances apart. Again, five positions within the same flow cell are chosen, and the separation between the two disks varies for each position: 200, 400, 600, 800, or 1000 μm apart. For K401 and Ncd experiments, the disks are illuminated for 30 frames, for Kif11 experiments, the disks are illuminated for 60 frames (frames are every 15 seconds). Then, a bar ≈ 5 μm wide connecting the disks is illuminated to merge the asters.

**Figure S2:**
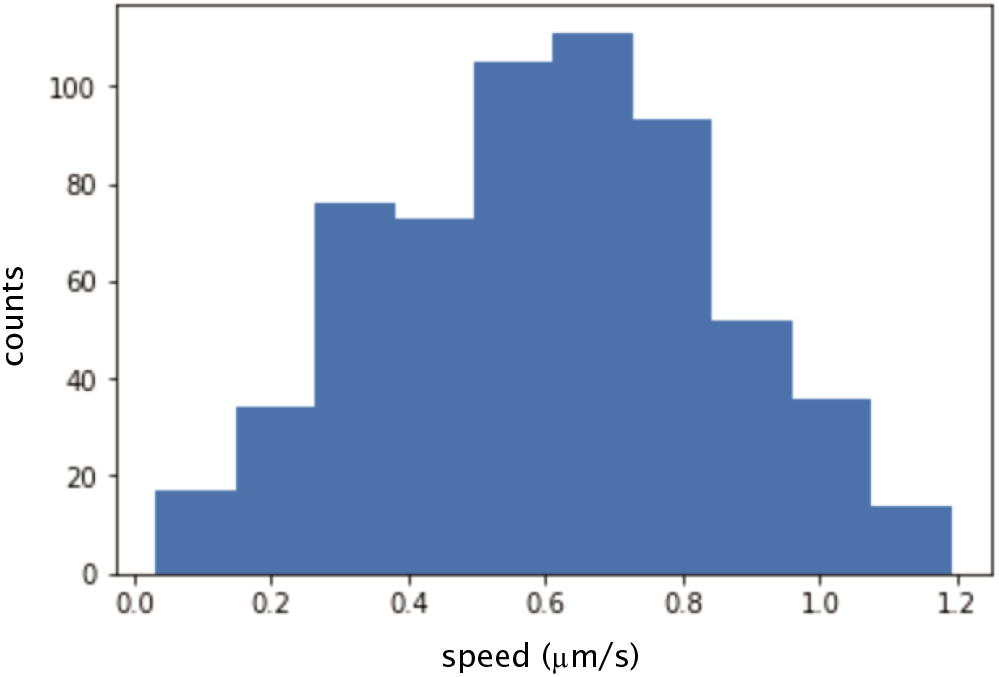
Histogram of calculated instantaneous speed of microtubules glided by K401 motors. The mean speed is ≈ 600 nm/s.

### 9.9 Gliding Assay

Motor speeds were determined by gliding assay. Glass slides and coverslips were Helmanex, ethanol, and KOH cleaned. Flow cells with ≈ 10 μL volume were created with double sided sticky tape, and rinsed with M2B buffer. Then, anti-GFP antibody was applied and incubated for 10 minutes. The flow cell was then rinsed with M2B and then mVenus labeled motor proteins (at ~ 5 nM in M2B) were flowed in and incubated for 10 minutes. The flow cell was rinsed with M2B to remove unbound motors and microtubules (in M2B with 3 mM ATP and 1 mM DTT) were flowed in. Microtubules were then imaged using total internal reflection fluorescence (TIRF) microscopy at a rate of one frame per second. Individual microtubules were tracked using custom written python code to determine their speed. The mean microtubule speed (excluding those that were not moving) was determined as the motor speed. Figure S2 shows the histogram of speeds obtained for K401 motors purified from SF9 cells.

### 9.10 Disordered Aster Core

We observe that the asters we create have centers that are very dense with motors and microtubules. By flourescence microscopy, we do not observe organized aster arms in this region and hypothesized that the microtubules are disordered in this region. To assess the extent of microtubule organization in our asters, we imaged asters with a polarized light microscope (Pol-Scope). This microscope utilizes polarized white light to image birefringent substances. Microtubules are birefringent due to their aspect ratio; they interact differently with light polarized parallel to their long axes compared to light polarized perpendicular to their long axes. Thus, the Pol-Scope allows determination of the alignment of microtubules, but not their plus/minus end polarity [38]. When imaged with a Pol-Scope, the arms of our asters are bright, indicating high alignment, and their azimuthal angle confirms that they are radially symmetric around the center (Fig. S3). The center of the aster is dark, which we interpret to mean that this region is disordered. It is possible that the microtubules in the center could be aligned pointing in the z direction, which could also result in the center being dark. A disordered center may be a result from steric hindrance due to a high density of microtubules in that region, which prevents the motors from aligning the microtubules. Due to the disorder in the aster center, we exclude this region from our theoretical analysis.

**Figure S3:**
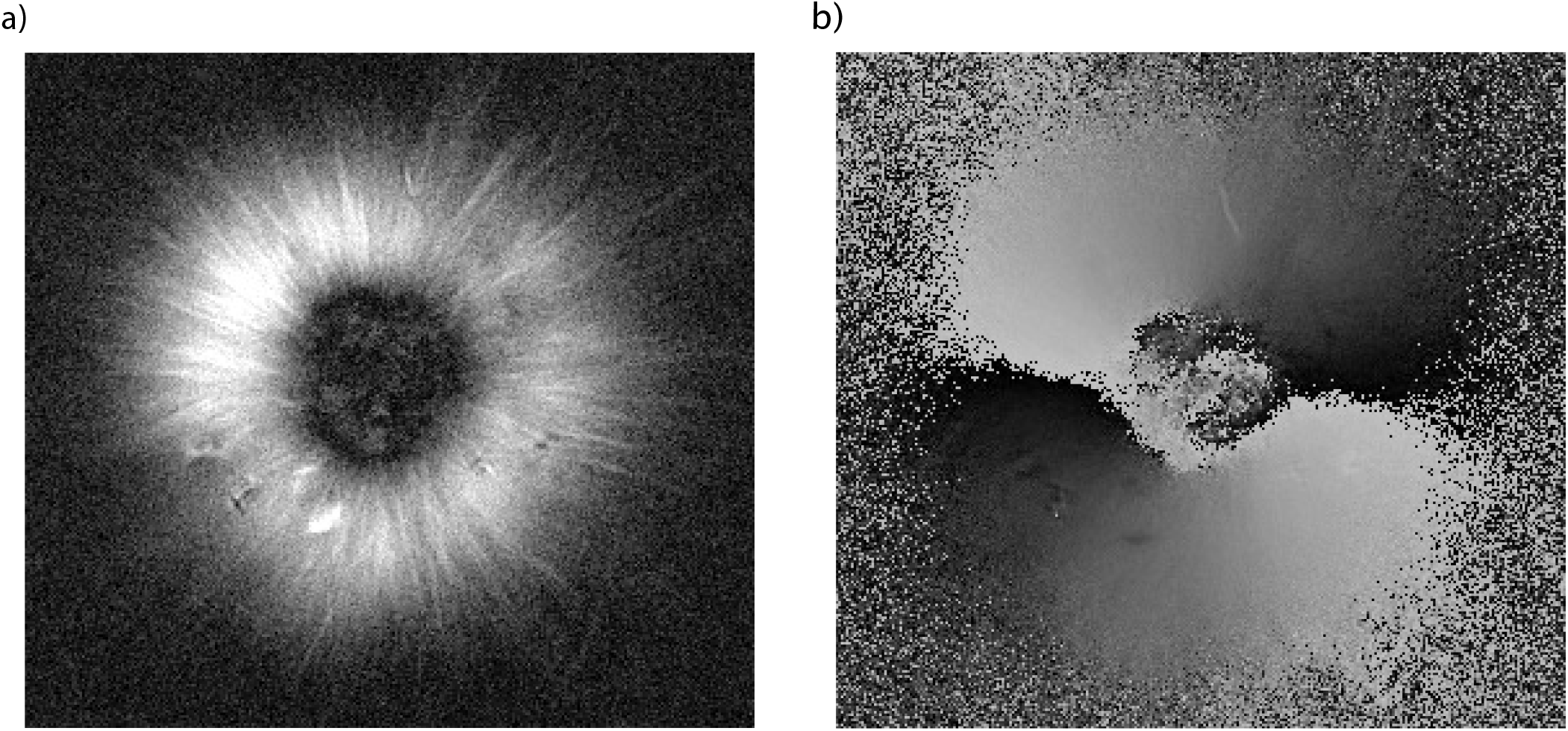
Aster centers are disordered and the arms are aligned radially. a) Retardance image of an aster taken with a Pol-Scope. The arms of the aster result from their alignment and the magnitude of retardance is proportional to the number of microtubules in a bundle. The dark center indicates no alignment of microtubules in that region. b) Azimuthal angle of microtubule alignment in an aster. Black is 0° and white is 180°.

### 9.11 Aster Size

We formed asters of various sizes using cylindrical illumination regions, ranging in diameter from 50 – 600 μm. Fig. S4 shows representative images of the microtubule fluorescence of asters formed with each motor and each excitation diameter. The yellow circles are by-hand determination of the outer boundary of the aster. In order to measure the size of asters in a more systematic way, we used the measured microtubule distribution, as shown in Fig. 1(C). Outside of the central core region of the aster, microtubule fluorescence decreases monotonically before rising again to the background level outside of the activation region. We chose to define the outer radius of the aster as the radius at which the median microtubule fluorescence is twice the background fluorescence. This metric agrees well with a visual inspection of the asters.

### 9.12 Merger Analysis

To measure contractile speeds in a pseudo one-dimensional network, we performed aster merger experiments. In these experiments, two asters were formed with 50 μm diameter excitation disks, at varying initial separations (either 200 μm, 400 μm, 600 μm, 800 μm, or 1000 μm). The two disks were illuminated for 30 frames or 60 frames for experiments with Kif11 (15 seconds between frames). After 30 or 60 frames, a thin rectangle was illuminated between the two asters. This caused the formation of a network which contracted, pulling the two asters together. An image of two asters connected in this way is shown in Fig. 3.

We used two methods to measure contractile rates. One was using optical flow to measure speeds throughout the network, as shown in Fig. 3(B). The open source package, Open CV was used for the optical flow measurements. We used a dense optical flow measurement using cv.CalcOpticalFlowFarneback, which calculates optical flow speeds using the Gunnar Farneback algorithm.

**Figure S4:**
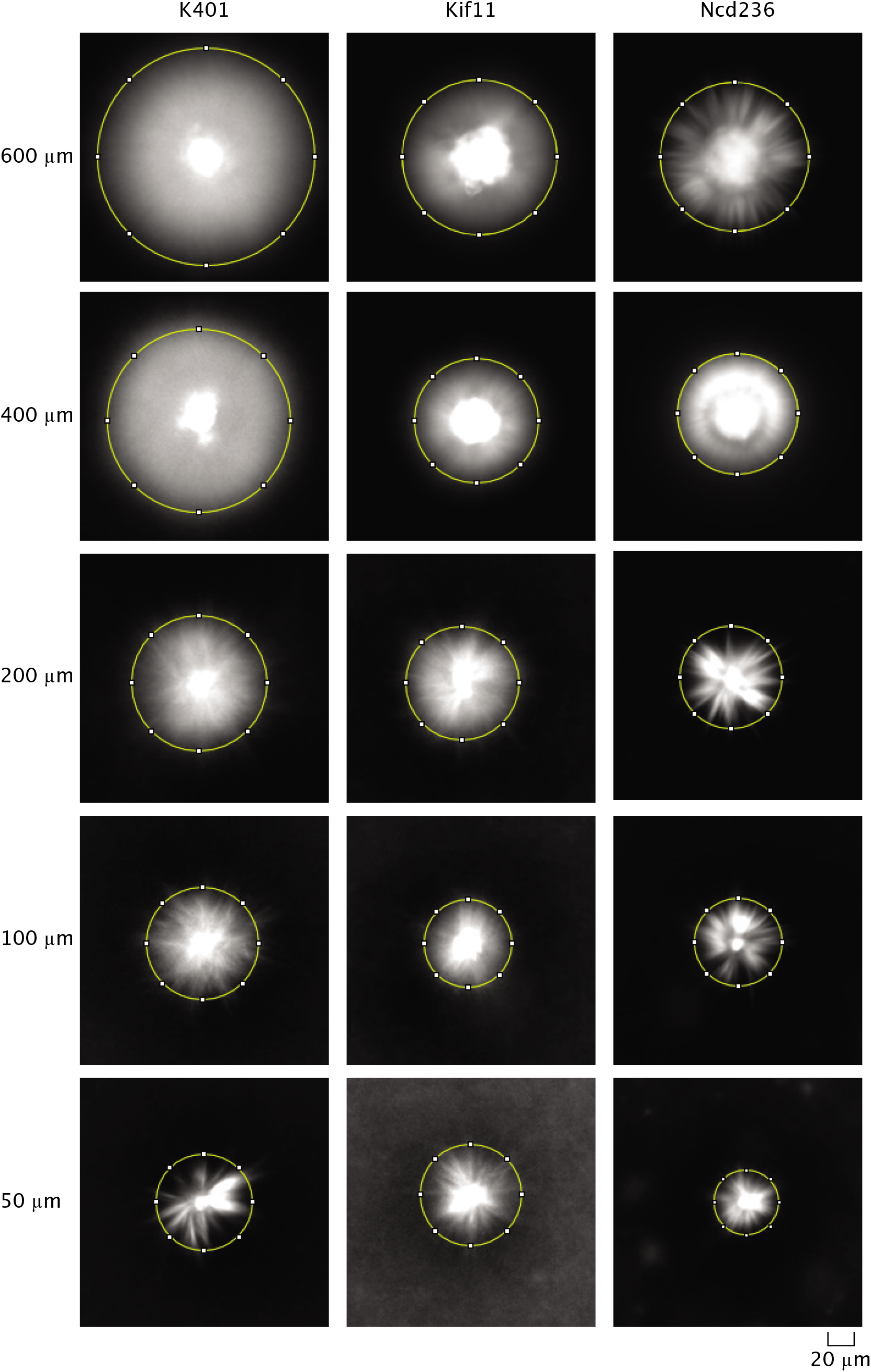
Example images of microtubule fluorescence of asters made with each motor used and each excitation diameter. The yellow circle indicates a visual measurement size of the aster.

**Figure S5:**
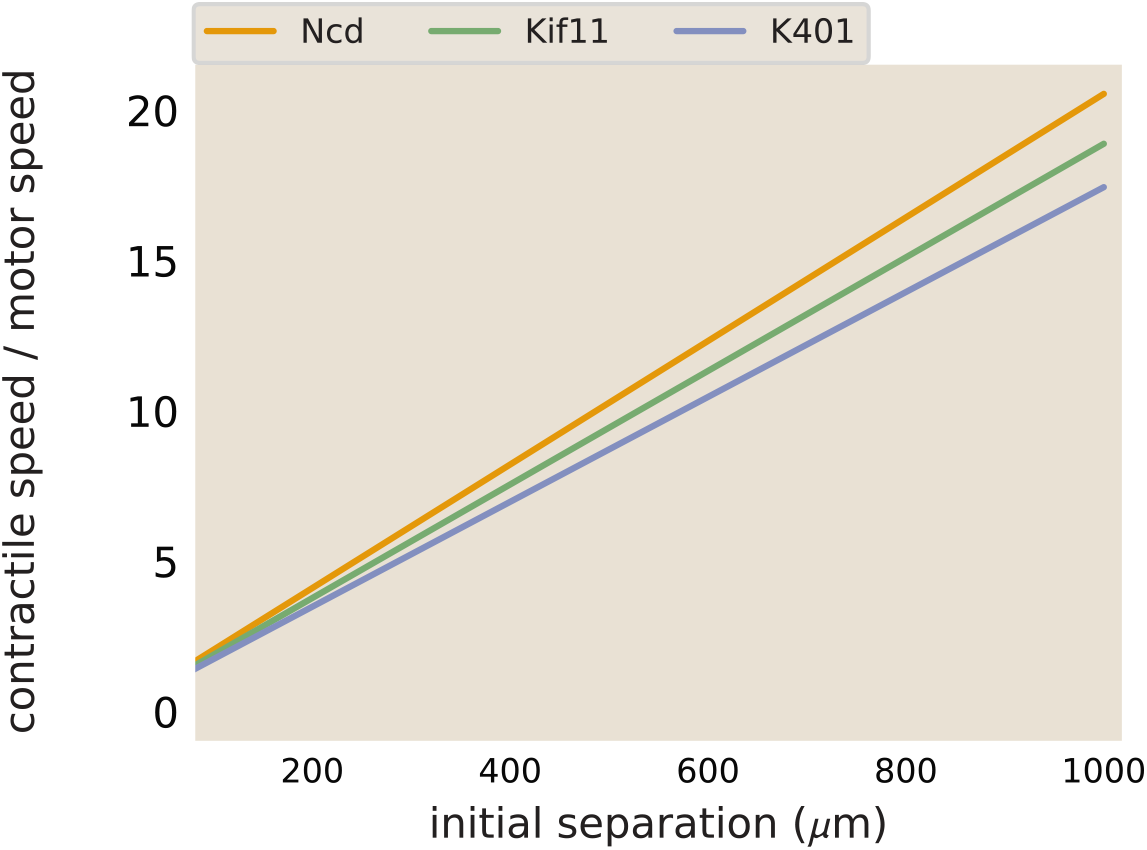
Merger speeds depend on motor stepping speed. The best fit lines from the contraction rate measured in aster mergers are plotted, normalized by the measured motor speed.

To calculate the maximum contractile rate, we tracked the positions of the asters over time. The microtubule fluorescence in the region containing the asters and network (pixels 450 – 650 in the y direction) was summed across the y dimension to make a one-dimensional line trace of fluorescence S6A. This trace was smoothed using a butter filter to reduce the noise in the fluorescence intensity. An example smoothed line trace is shown in Fig. S6A. The asters were then identified as the large peaks in fluorescence intensity using scipy.signal.find_peaks. The identified peaks are shown as red dots in Fig. S6A. The corresponding image that the line trace is from is shown in Fig. S6B with a 50 x 50 pixel box around the identified aster colored in black. From the identified aster coordinates, the distance between the asters was calculated and the difference between distances in successive frames was used to calculate the merger speed. The reported speed is the speed of a single aster. Thus, the speed calculated from the change in distance between asters is divided by two, assuming the asters are moving at equal speed.

Example traces of calculated aster speeds over time after the beginning of illuminating the bar region is shown in Fig. S6c. The speed of the asters increases while the network connecting them forms, peaks, and then decreases as the asters near each other. The peak speed is reported as the maximum aster speed in the main text. These maximum aster speeds as a function of initial separation were fit to a line using scipy.optimize.curve_fit to determine the slope of the increase in speed with separation. The best fit lines determined with this method are plotted along with the data for Ncd and Kif11 in Fig. 3c. The fit slope for Ncd was 0.0024 s^-1^ and 0.0013 s^-1^ for Kif11. The ratio (Ncd/Kif11) of these slops is ≈ 1.8. In comparison, the ratio between measured motor speeds (115 nm/s for Ncd and 70 nm/s for Kif11) is ≈ 1.6. The slope calculated for K401 was ≈ 0.0043 s^-1^. Thus, the ratio of slopes of aster speeds versus separation is in good agreement with the ratio of motor speeds, suggesting that the motor speed sets this slope. To illustrate this point, the best fit lines shown in Fig. 3c are plotted again, with the slopes divided by the single motor speed. In this way, the three lines now overlap, as shown in Fig. S5.

### 9.13 Model of Network Contraction

In our aster merger experiments, we observed a linear relationship between distance from the center of the network and aster speed. This is seen in the plot of x-velocity vs position in Fig. 3(B) and also in the plot of maximum aster speed versus initial separation in Fig. 3(C). Juniper et al. observed a similar relationship between position from the center of the network and contraction speed during aster formation in [39]. We interpret this linear trend to mean that the contractile network can be thought of as a series of independently contracting units. Going farther away from the center of the network adds more contractile units, generating the observed speed increase. One question that remained, however, is what sets the slope of the increase of the measured contractile speed. We hypothesize that the single motor speed is one factor. This results in an equation for the rate of contraction in a network to scale as

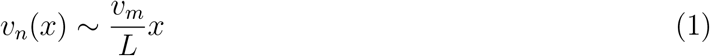

where v_*n*_ is the network contractile rate, v_*m*_ is the motor speed, L is a characteristic length scale, and x is the position from the center of the network. Our observed relationship that the ratio of the slope of speed increase matched the ratio of single motor speeds (as discussed in the main text) supports this hypothesis. Further, we can estimate that L ≈ 50 – 100 μm, indicating that the characteristic length scale is larger than a single microtubule.

### 9.14 Motor Competition

To test the effects of competing motors, we performed experiments with two motors at a time. The reaction mixture was the same as other experiments, with the two motors added at varying concentrations. With our light activation, we were able to not only form motor homodimers (K401- K401, for example), but also heterodimers that consist of a sets of motor heads that differ (e.g. K401-Ncd). These motor heterodimers set up a direct competition between the motor heads, with one end set of motor heads walking towards microtubule plus-ends and the other set walking towards microtubule minus-ends. In order to create an aster, it is thought that one motor must ‘win’ to polarity sort the microtubules such that either the minus ends or the plus ends are brought together in the center.

We found that depending on the relative concentrations of the two motors, one of several outcomes would occur: aster with the plus-end directed motor in the center, aster with the minus-end directed motor in the center, or failed contraction. In the stalled or failed contraction case, as is the case with K401-Ncd or Kif11-Ncd, bundles of microtubules are formed that do not lead to the reorganization necessary to form an aster. In this case, we see that the fluorescence of mCherry (Kif11) and YFP (Ncd) are about equal and uniform across the activated region (Fig. S7A). In contrast, with excess Ncd dimers, an aster is formed where Ncd accumulates in the center and Kif11 is more prevalent in the aster arms. This can be seen in Fig. S7B, where the ratio of Kif11:Ncd fluorescence increases moving outward from the aster center.

**Figure S6:**
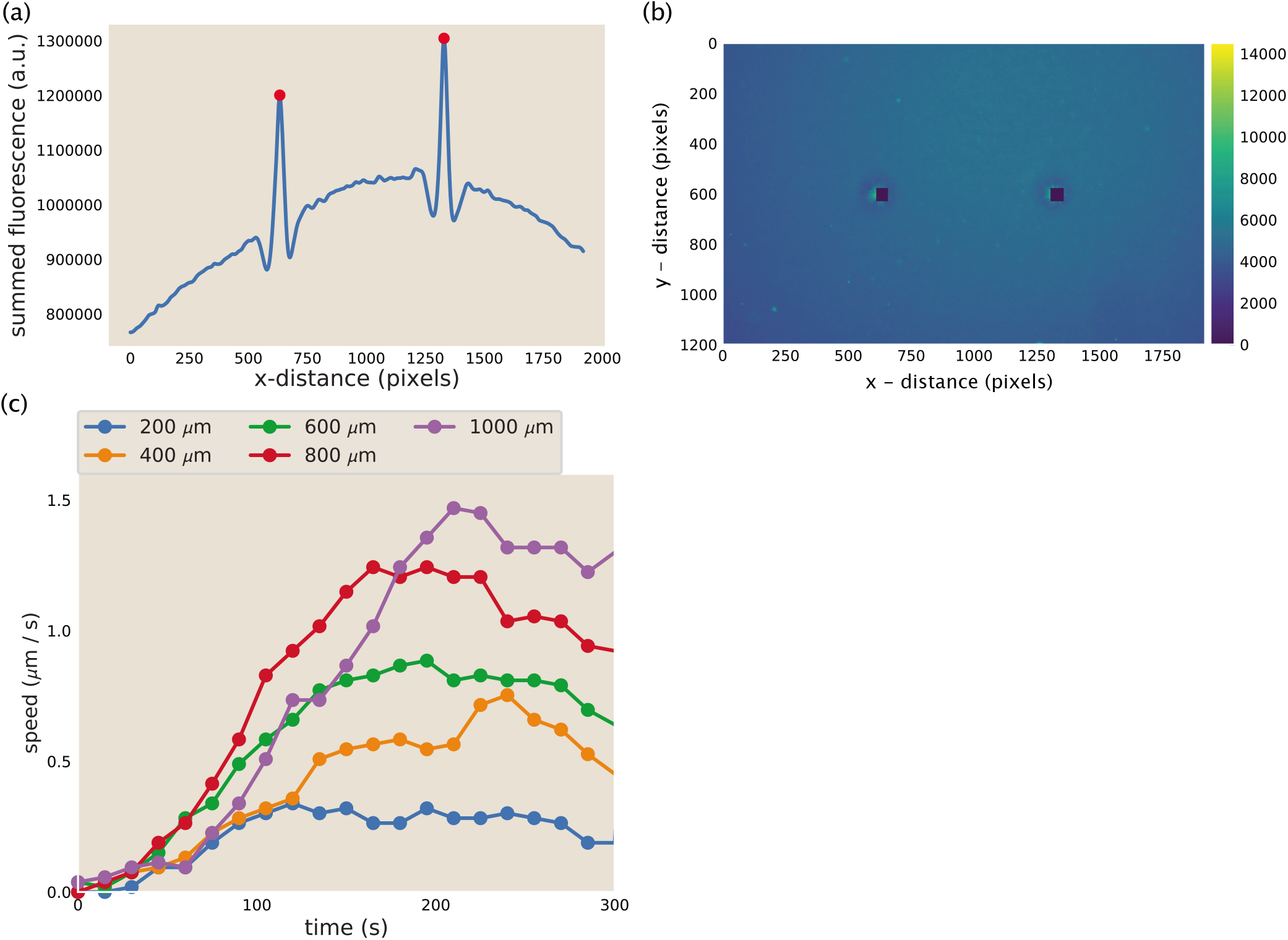
Aster identification and speed calculation in aster mergers. (a) Example y-summed microtubule fluorescence from an image during aster merger. The peaks represent the asters and the red dots are the position of the asters as identified by the code. (b) Image corresponding to the fluorescence plotted in (a). The blacked out squares are the location of the asters identified from the peaks in (a). (c) Measured aster speed versus time during aster merger. Each dot is a single measured speed and the colors represent the initial separation of the asters.

**Figure S7:**
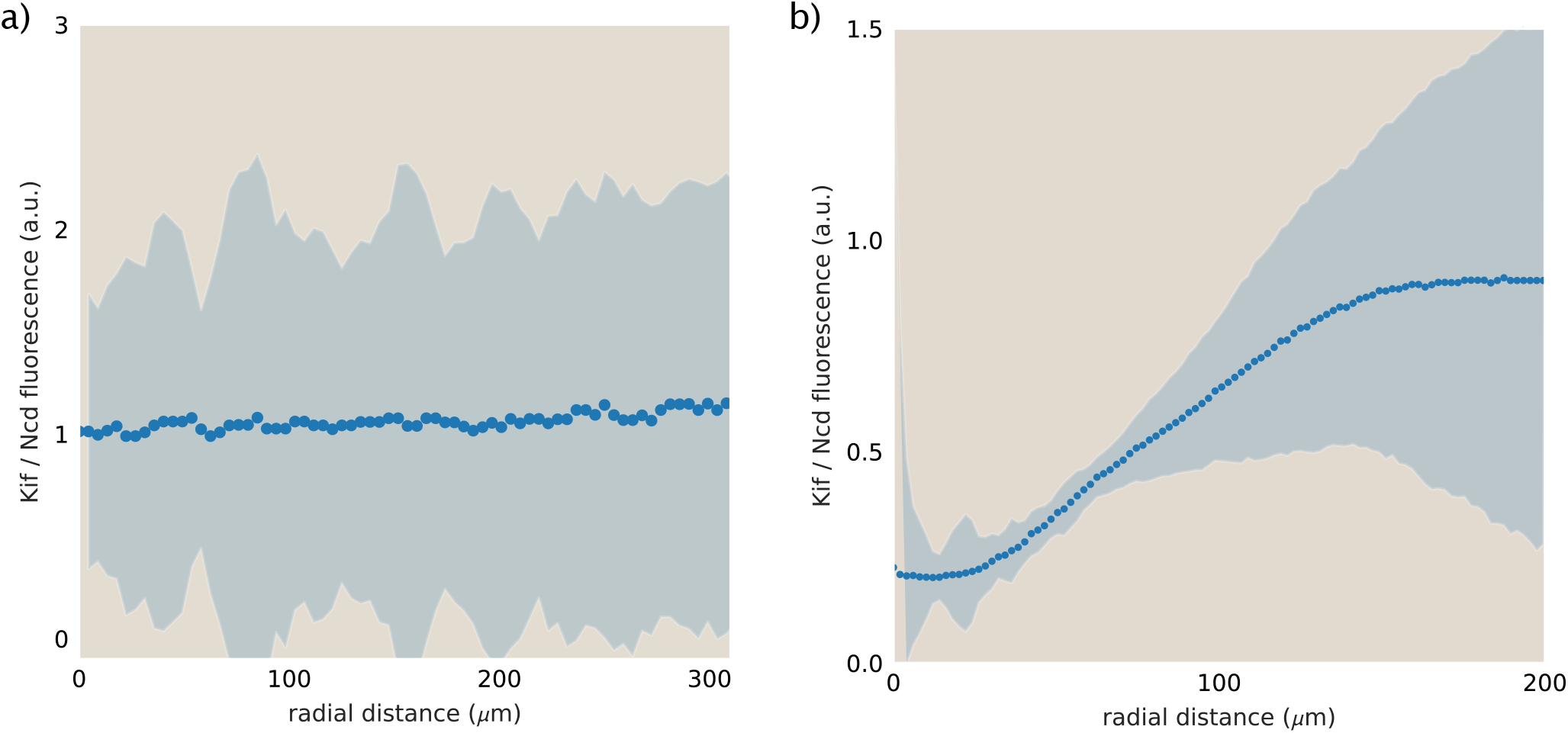
Distributions of Kif11 and Ncd in competition experiments. Ratio of Kif11:Ncd fluorescence from an experiment with Kif11-Ncd heterodimers (a) and Ncd-Ncd and Kif-Ncd (b). The shading represents one standard deviation.

### 9.15 Motor distributions

#### 9.15.1 Model formulation

To predict the spatial distribution of motors in aster structures, we model the dynamic steady-state of free (‘f’) and bound (‘b’) motor concentrations via

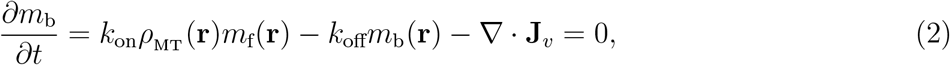

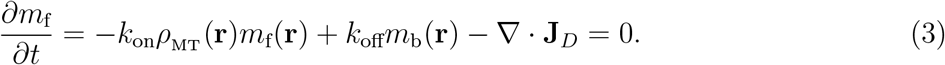

Here, *k*_on_ and *k*_off_ are the motor binding and unbinding rates, respectively, *ρ*^MT^(**r**) is the spatially varying microtubule concentration at steady state (measured as μM tubulin), **J**_*v*_ is the advective flux of bound motors, and **J**_*D*_ is the diffusive flux of free motors. Our modeling approach is similar to that used by Nédélec, *et al*. [22] with the main difference being in the handling of *ρ*_MT_(**r**). Namely, they imposed a particular functional form on this distribution (*ρ*_MT_(**r**) = 1/|**r**|^*d*-1^ with *d* as the spatial dimension) based on an idealized representation of microtubule organization in an aster, whereas in our treatment *ρ*_MT_(**r**) stands for the experimentally measured microtubule profiles which cannot be captured through an analogous idealization.

If the free motors have a diffusion coefficient *D*, then, in the radially symmetric setting considered in our modeling, the diffusive flux will be given by

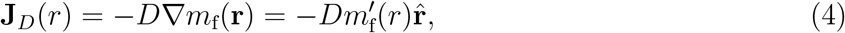

where 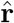 is an outward-pointing unit radial vector. And if *v* is the advection speed of bound motors, then the advective motor flux on radially organized microtubules will be

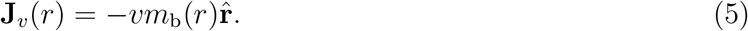

Here, we are implicitly assuming that motors constantly walk when bound, ignoring the fact that they can stall upon reaching a microtubule end. We discuss the impact of this effect later in Appendix section 9.15.5.

At steady state, the net flux of motors at any radial distance *r* should be zero (**J**_*D*_ (*r*) + **J**_*v*_ (*r*) = 0), which implies a general relation between the profiles of free and bound motors, namely,

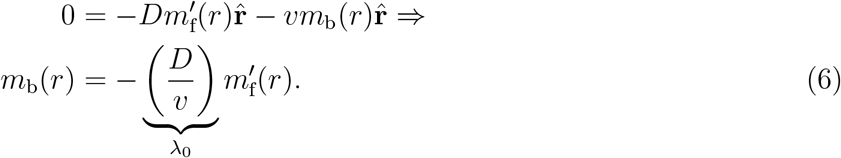

Above we introduced λ_0_ as a length scale parameter that can be interpreted as the distance which is traveled by free and bound motors at similar time scales, i.e., diffusion time scale 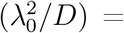 advection time scale (λ_0_/*v*). Note also that the ‘-’ sign at the right-hand side indicates that the free motor population should necessarily have a decaying radial profile 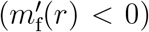 which is intuitive since at steady state the outward diffusion needs to counteract the inward advection.

To make further analytical progress, we will assume that motor binding and unbinding events are locally equilibrated [25]. This assumption is valid if motor transport is sufficiently slow compared with binding/unbinding reactions. We will justify this quasi-equilibrium condition for the motors used in our study at the end of the section. It follows from this condition that

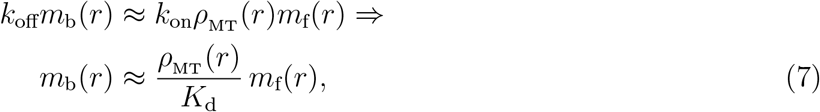

where *K*_d_ = *k*_off_/*k*_on_ is the dissociation constant. Since the experimental readout reflects the *total* motor concentration (*m*_tot_ = *m*_f_ + *m*_b_), we use our results (Eq. 6 and Eq. 7) to link *m*_tot_ (*r*) with the microtubule profile *ρ*_MT_ (*r*). Specifically, using Eq. 7 we find

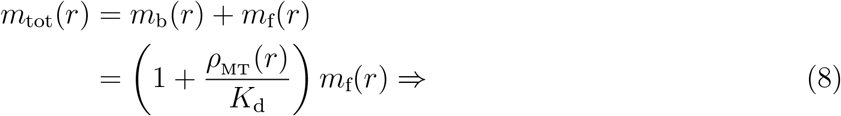

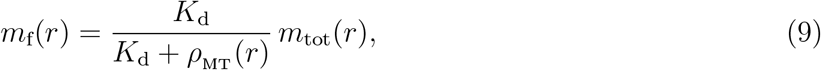

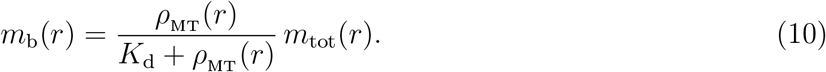

Next, substituting the above expressions for *m*_f_ and *m*_b_ into Eq. 6 and simplifying, we relate the motor and microtubule profiles, namely,

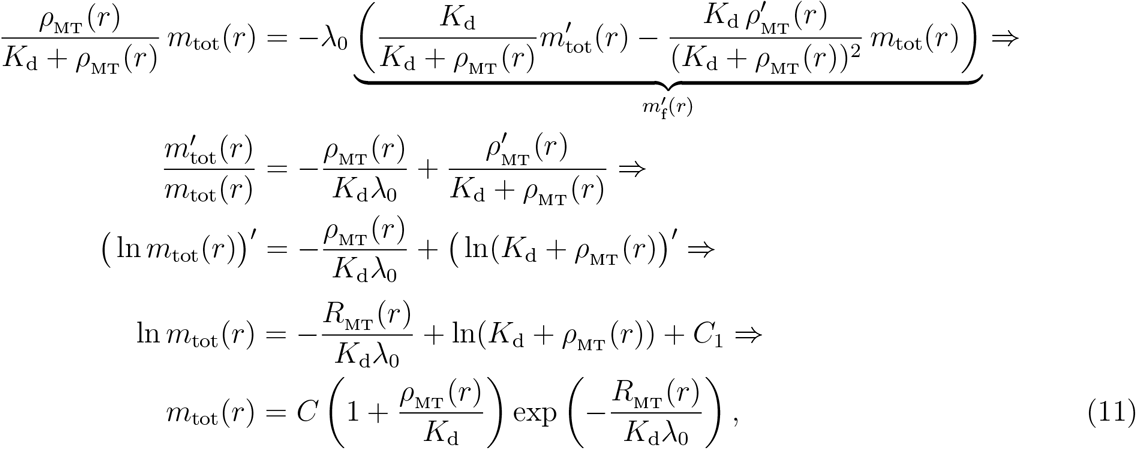

where *R*_MT_ (*r*) = ∫ *ρ*_MT_ (*r*) d*r* is the integrated microtubule concentration, and *C* = *K*_d_ *e*^*C*_1_^ is a positive constant. The presence of the multiplicative constant C is a consequence of the fact that the two equations used for deriving our result (Eq. 6 and Eq. 7) specify the *ratios* of motor populations. Therefore, the result in Eq. 11 predicts the relative level of the total motor concentration, given the two effective model parameters (*K*_d_ and λ_0_), which we infer in our fitting procedure.

Note that the two variable factors on the right-hand side of Eq. 11 have qualitatively different structures. The first one is local and depends only on the dissociation constant (an equilibrium parameter), while the second term involves an integrated (hence, non-local) microtubule density term and λ_0_ = *D/v* which depends on the advection speed *v* (a non-equilibrium parameter). As anticipated, in the limit of vanishingly slow advection (*v* → 0 or, λ_0_ → ∞) the second factor becomes 1 and an equilibrium result is recovered.

##### Connections to related works

Before proceeding further into our analysis, we briefly compare the expression for the motor distribution (Eq. 11) with analogous results in the literature. Specifically, Nédélec, *et al.* [22] studied quasi-two-dimensional asters and in their modeling treated microtubules as very long filaments, all converging at the aster center. This setting implied ~ 1/r scaling of the microtubule concentration. With this scaling, the integrated microtubule concentration in our framework becomes 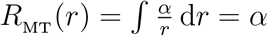 ln *r* where *α* is a constant. Substituting this form into the exponential term in Eq. 11, we find 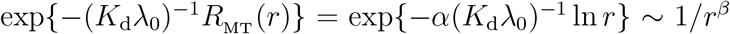, where *β* = *α*(*K*_d_λ_0_)^-1^. It then follows from Eq. 11 and the scaling *ρ*_MT_(*r*) ~ 1/*r* that the motor concentration is a sum of two decaying power laws (namely, ~ 1/*r^β^* and ~ 1/*r*^*β*+1^) – the result obtained by Nédélec, *et al.* [22]. A more detailed calculation can be done to demonstrate that the exponent *β* matches exactly with the result derived in the earlier work, but for the purposes of our study we do not elaborate further on this comparison. We note that the experimentally measured microtubule profiles in asters (e.g., Fig. 2b or Fig. 3d) often have an inflection point and cannot be fitted to decaying power law functions (e.g., 1/r^2^ for 3D asters), which is why the idealized setting considered by Nédélec, *et al.* [22] cannot be applied to our system.

Another set of works [23, 24] also studied motor distributions in asters, but this time under the assumption of a uniform microtubule concentration (*ρ*_MT_(*r*) ~ constant). In such a setting, our framework predicts an exponentially decaying motor profile, because *R*_MT_ (*r*) = ∫ *ρ*_MT_ (*r*) d*r* ~ *ρ*_MT_*r* and thus, 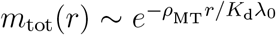. An exponential decay was also the prediction of Lee and Kardar [23], although in their treatment all motors were assumed to be in the bound state. The two distinct motor states were considered in the work by Sankararaman, *et al.* [24] who predicted an exponential decay of motor concentration modulated by a power-law tail. One can show, however, that when the decay length scale of motor concentration greatly exceeds the motor processivity (as in the case of asters which we generated), the prediction of Sankararaman, *et al.* [24] also reduces into a pure exponential decay, matching the prediction of our model. But since the assumption of a uniform microtubule profile is clearly violated in our system, these predictions are not applicable for us.

##### Validity of the quasi-equilibrium assumption

Earlier in the section, we assumed that motor binding and unbinding reactions were locally equilibrated, from which Eq. 7 followed. Looking at the governing equation of bound motor dynamics (Eq. 2), we can see that this assumption will hold true if *k*_off_*m*_b_(*r*) » |∇ · **J**_*v*_|. Substituting the expression of advective flux (Eq. 5) and recalling that in three dimensions the divergence of a radial vector 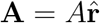 takes the form *r*^−2^*∂_r_* (*r*^2^*A*), we rewrite the quasi-equilibrium condition as

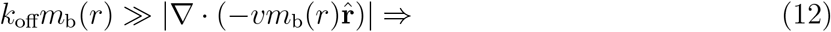

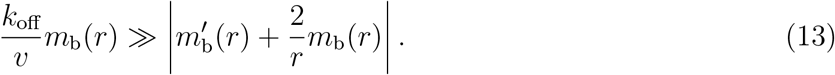

Now, many of the motor profiles can be approximated reasonably well by an exponentially decaying function (see Fig. S10 for a collection of experimental profiles). This suggests an empirical functional form *m*_b_(*r*) ~ *e*^-*r*/λ^ for the concentration of bound motors, where λ is the decay length scale (note that the constant saturation level contributes to the *free* motor population). This functional form implies that 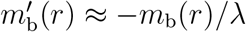, which, upon substituting into Eq. 13, leads to

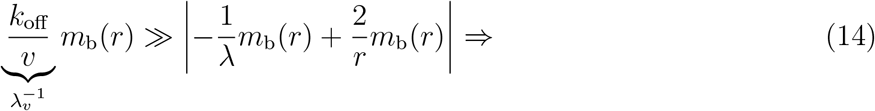

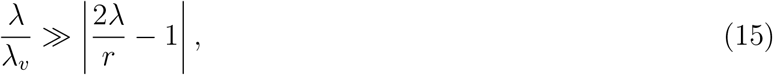

where λ_*v*_ = *v/k*_off_ is introduced as the motor processivity (distance traveled before unbinding). The processivities (λ_*v*_) of the three different kinesins used in our study, together with the observed ranges of decay length scales (*λ*) of corresponding motor profiles are listed in Table S2. As can be seen, in all cases the ratio λ/λ_*v*_ is much greater than one, verifying the intuitive expectation that the length scales of aster structures are much greater than the single run lengths of motors.

It is obvious from the presence of the *r*^-1^ term on the right-hand side of Eq. 15 that the condition can only be satisfied past a certain radius, since *r*^-1^ becomes very large when *r* approaches zero. This threshold radius (*r*\*) is set by *r*\* ~ 2λ_*v*_, where the two sides of Eq. 15 become comparable. The threshold radial distance that we choose to isolate the core is at least 5 – 10 μm for the asters of our study (see the lower *x*-limits in the profiles of Fig. S10) which exceed *r*\* at least a few times. This suggests that Eq. 15 is valid, justifying our use of the quasi-equilibrium assumption for modeling the motor distribution.

**Table S2:** Processivities of motors, decay length scales of motor profiles, and corresponding ratios of these two length scales. Step size of ≈ 10 nm corresponding to the length of a tubulin dimer was used for estimating the motor processivities in μm units. For Ncd motors, the upper limit in processivity corresponds to that of oligomeric motor assemblies. Estimates for the decay length scales *λ* were made based on the motor profiles in Fig. S10.

| motor | processivity (steps) | processivity, $\lambda_v$ ($\mu\text{m}$) | decay length scale, $\lambda$ ($\mu\text{m}$) | $\lambda/\lambda_v$ |
| --- | --- | --- | --- | --- |
| K401 | $\approx 100$ | 1 | 10–40 | 10–40 |
| Ncd236 | $\approx 1-100$ | $10^{-2} - 1$ | 5–20 | 5–2000 |
| Kif11(513) | $\approx 10$ | $10^{-1}$ | 10–20 | 100–200 |

#### 9.15.2 Extraction of concentration profiles from raw images

In this section, we describe our approach for extracting the radial profiles of motor and microtubule concentrations from raw fluorescence images.

##### Fluorescence normalization and calibration

When taking images with a microscope, several sources contribute to the detected pixel intensities: the camera offset, autofluorescence from the energy mix, and fluorescence coming from the tagged proteins (tubulin or motors). In addition, due to the uneven illumination of the field of view, the same protein concentration may correspond to different intensities in the raw image.

We begin the processing of raw images by first correcting for the uneven illumination. For microtubule images, we use the first movie frames as references with a uniform tubulin concentration in order to obtain an intensity normalization matrix. Each pixel intensity of the final image frame is then rescaled by the corresponding normalization factor.

Although the motor concentration is also initially uniform, the light activation region in the first frame appears photobleached, making it unsuitable for the construction of a normalization matrix. Instead, we obtain this matrix from the final frame, after masking out the neighboring region of the aster, outside of which the nonuniformity of the fluorescence serves as a proxy for uneven illumination. Intensity normalization factors inside the masked out circular region are obtained through a biquadratic interpolation scheme. The steps leading to a normalized motor image are depicted in Fig. S8a.

After fluorescence normalization, we convert intensities into units of protein concentration using calibration factors estimated from images of samples with known protein contents. For K401 and Kif11 motors, we use the conversion 1000 intensity units → 815 nM motor dimer. For Ncd dimers, which have fluorescent tags on both iLid and Micro units, we use the 1000 intensity units → 407 nM conversion. In all three cases, 200 ms exposure time is used in the imaging. For tubulin, we make a rough estimate that after spinning the energy mix with tubulin, around 1 μM of tubulin remain, all of which polymerize into GMP-CPP stabilized microtubules. This leads to the calibration of 360 intensity units → 1 μM tubulin (100 ms exposure time).

##### Aster center identification

In the next step of the profile extraction pipeline, we crop out the aster region from the normalized image and identify the aster center in an automated fashion. In particular, we divide the aster into 16 equal wedges, calculate the radial profile of motors within each wedge, and define the aster center as the position that yields the minimum variability between the motor profiles extracted from the different wedges. Having identified the center, a mean radial profile for the aster is defined as the average of the 16 wedge profiles (Fig. S8b).

##### Inner and outer boundary determination

Since our modeling framework applies to regions of the aster where the microtubules are ordered, we consider the concentration profiles in a limited radial range for the model fitting procedure. As we do not have a PolScope image for every aster to precisely identify the disordered core region, we prescribe a lower threshold on the radial range by identifying the position of the fastest intensity drop and adding to it a buffer interval (equal to 15% of the outer radius) to ensure that the region of transitioning from the disordered core into the ordered aster ‘arms’ is not included (Fig. S8c, top panel). As for the outer boundary, we set it as the radial position where the tubulin concentration exceeds its background value by a factor of two (Fig. S8c, bottom panel).

We refer the reader to Supplementary code files for more details on the profile extraction procedure.

**Figure S8:**
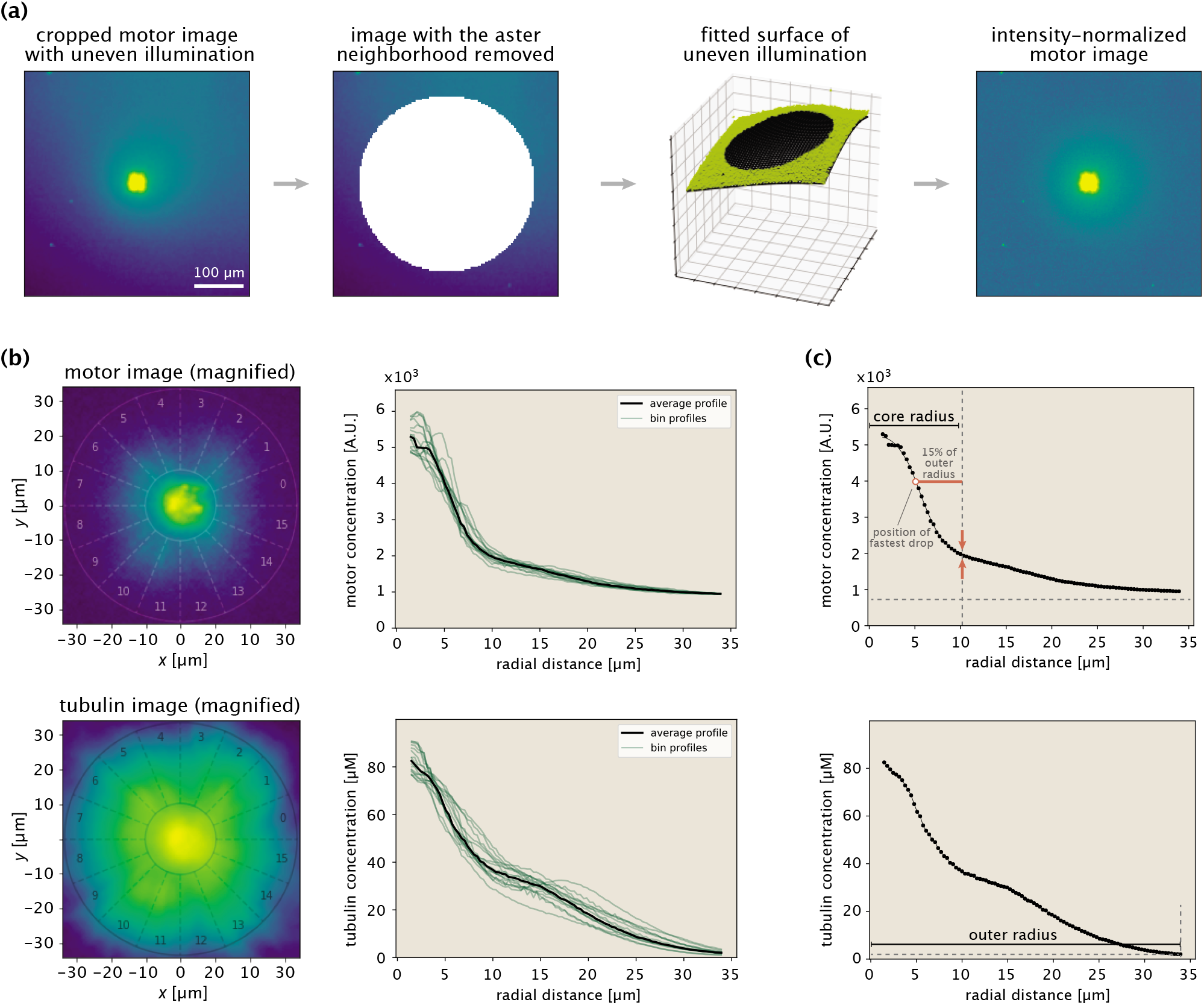
Procedure for extracting protein concentration profiles demonstrated on an example aster. (a) Steps taken in normalizing the fluorescence of motor images. The immediate aster region is shown with a saturated color to make it possible to see the nonuniform background fluorescence. (b) Aster center identification and extraction of radial concentration profiles. The numbers indicate the wedges at different angular positions. The two circles in the images indicate the inner and outer bounds. (c) Determination of inner and outer bounds based on the motor and tubulin profiles, respectively.

#### 9.15.3 Model fitting

Here we provide the details of fitting the expression we derived for the motor distribution (Eq. 11) to the profiles extracted from aster images. Since smaller asters are typically irregular and hence, do not meet the polar organization and radial symmetry assumptions of the model, we constrain the fitting procedure to larger asters formed in experiments with a minimum light illumination disk diameter of 200 μm.

The different aspects of the fitting procedure are demonstrated in Fig. S9. Extracting the average tubulin and motor profiles, we fit our model to the motor profile and obtain the optimal values of the effective parameters *K*_d_ and λ_0_. With the exception of a few cases, the optimal pair (*K*_d_,λ_0_) corresponds to a distinct peak in the residual landscape (Fig. S9c, note the logarithmic scale of the colorbar), suggesting that the parameters are well-defined. The fit to the motor data for the example aster is shown in Fig. S9d.

**Figure S9:**
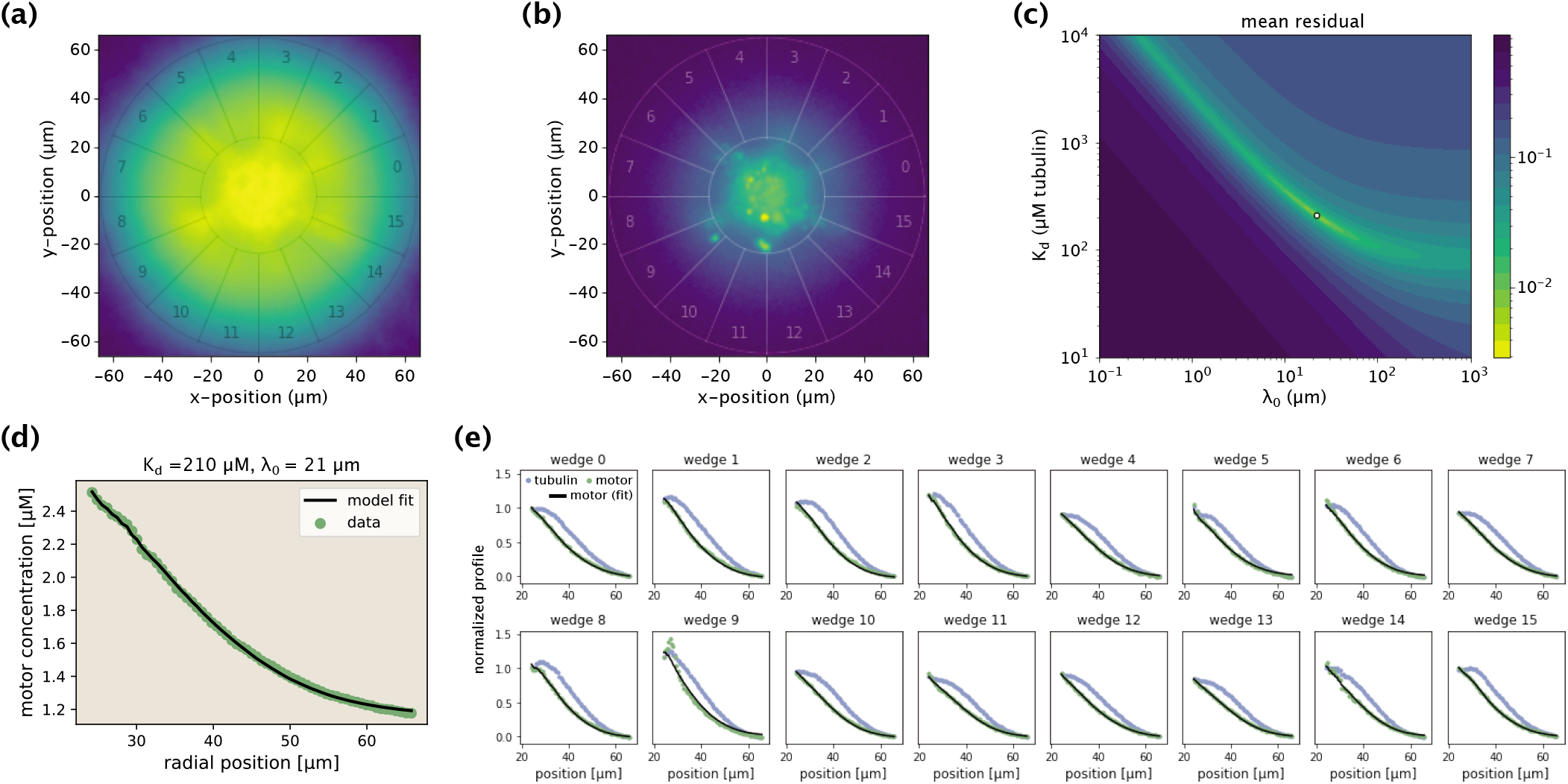
Demonstration of the model fitting procedure for average as well as separate wedge profiles. (a,b) Fluorescence images of an example Kif11 aster in tubulin (a) and motor (b) channels. 16 different wedges are separated and numbered. (c) Landscape of fit residuals when varying the two effective parameters *K*_d_ and λ_0_. For each pair, an optimal scaling coefficient *C* is inferred before calculating the residual. The dot at the brightest spot stands for the optimal pair (or, the arrow indicates the location of the optimal pair in the landscape). (d) Average motor profile and the model fit, along with the inferred parameters. (e) Collection of fits to separate wedge profiles using the optimal (*K*_d_, λ_0_) pair inferred from the average profile.

As stated in the main text, we then use the data from the separate aster wedges to assess the quality of fit for each aster. Specifically, keeping the (*K*_d_, λ_0_) pair inferred from the average profile fixed, we fit the 16 separate wedge profiles by optimizing over the scaling coefficient *C* for each of the wedges, and use the model residuals to assess the fit quality. In the set shown in Fig. S9e, with the exception of wedge 9 which contains an aggregate near the core radius, fits to all other wedge profiles are of good quality, translating into a low fitting error reported in Fig. 2e of the main text.

Repeating this procedure for all other asters, we obtain the best fits to their motor profiles and the corresponding values of the optimal (*K*_d_, λ_0_) pairs. The collection of all average profiles, along with the best model fits and inferred parameters are shown in Fig. S10.

#### 9.15.4 Expected ratio of *K*_d_ values for K401 and Kif11 motors

Here we show the steps in estimating the expected ratio of *K*_d_ values for the motors K401 and Kif11. Recall the definition *K*_d_ = *k*_off_/*k*_on_. Knowing the motor processivities and speeds from Table 1, we calculate the off-rates as *k*_off_ = speed/processivity. This yields a ratio 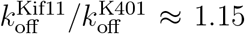. Then, using the reported on-rates in Valentine and Gilbert [26], we find the ratio of on-rates to be 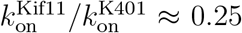. Taken together, these two results lead to our estimate for the ratio of *K*_d_ values reported in the main text, namely, 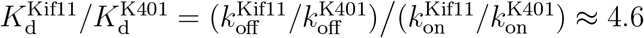.

**Figure S10:**
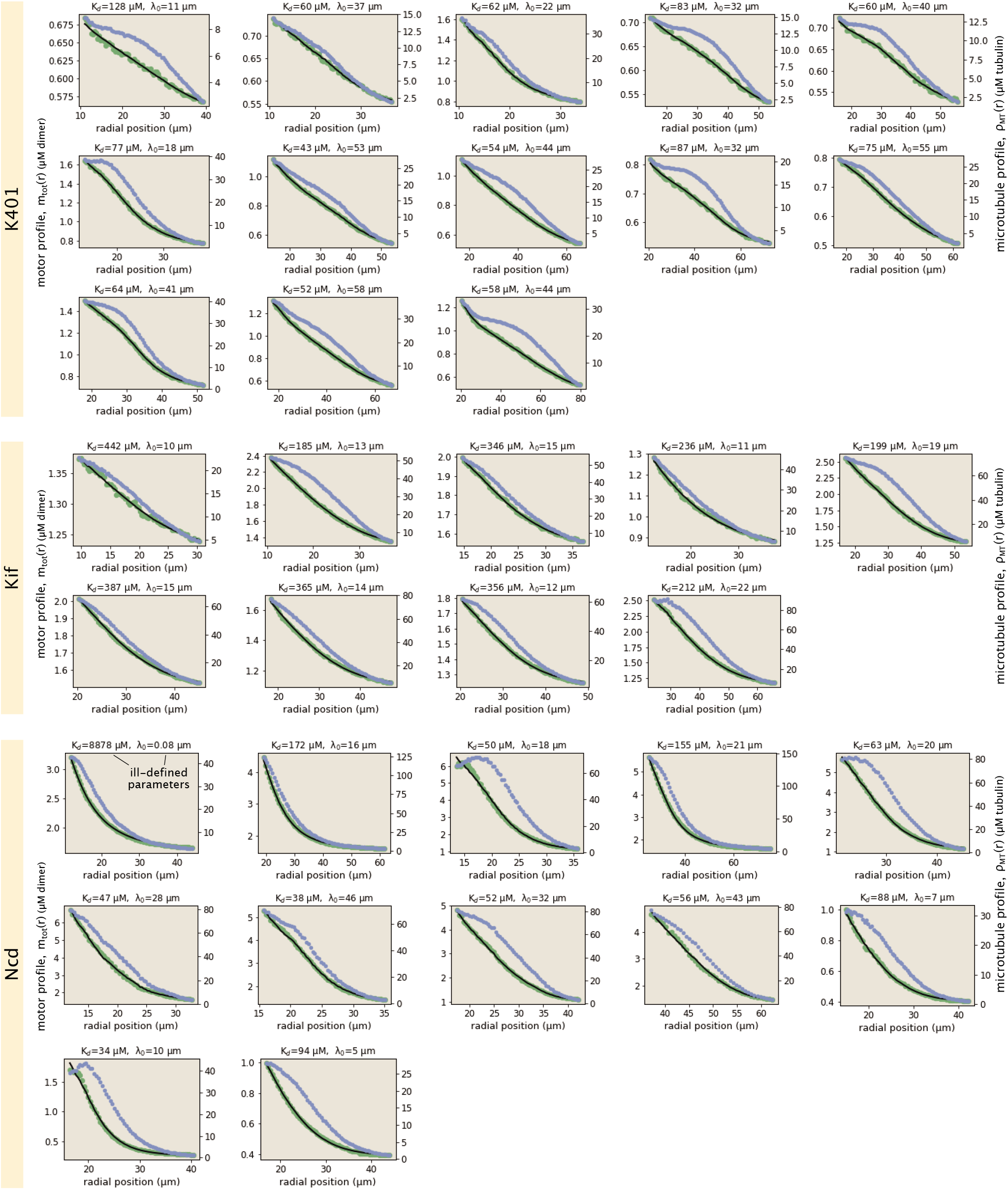
Collection of all fits to motor profiles. The green and blue dots represent the radially averaged motor and tubulin concentrations. The solid black lines represent the model fits.

#### 9.15.5 Accounting for finite MT lengths

Analysis of purified microtubule images shows that the median length of microtubules is ≈ 1.6 μm (Fig. S1). Taking the size of a tubulin dimer to be 8 nm, this length translates into the distance traveled in ≈ 200 motor steps, which is comparable to the processivity of K401 motors reported in Table 1 of the main text. Since motors stall when reaching microtubule ends, their effective advection speed will get reduced. Here, we account for this reduction and estimate its magnitude for the different motors used in our study.

Consider the schematic in Fig. S11 where a motor is shown advecting on a microtubule with length *L*. If the distance *x* between the motor and microtubule end at the moment of binding is less than the motor processivity λ_*v*_, then the motor will reach the end and stall for a time period 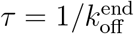 before unbinding. On the other hand, if *x* is greater than λ_*v*_, the motor will not stall while bound to the microtubule and hence, its effective speed will not be reduced.

**Figure S11:**
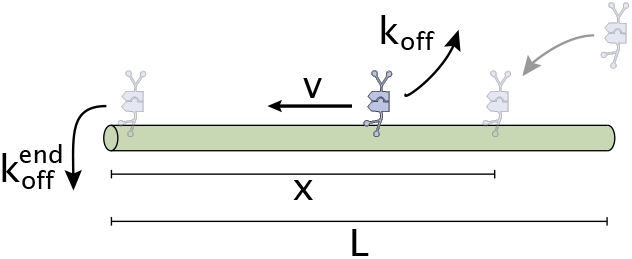
Schematic representation of initial motor binding, advection and stalling at the microtubule end.

Assuming that the location of initial binding is uniformly distributed in the [0, *L*] interval (hence, the chances of binding between *x* and *x* + d*x* is d*x*/*L*), we can calculate the effective speeds in the above two cases as

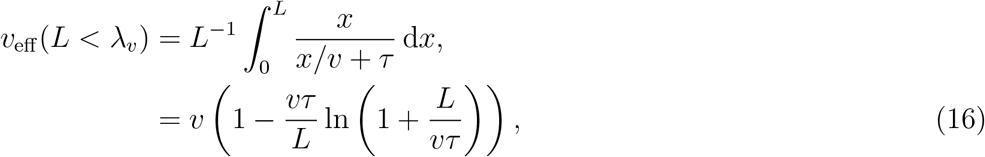

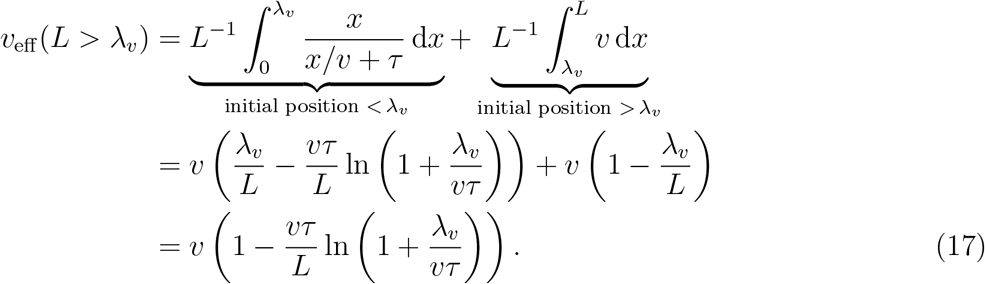

As can be seen, in both cases the effective speed is lower than the walking speed *v*. Now, if *p*(*L*) is the distribution of microtubule lengths, then the mean effective motor speed evaluated over the whole microtubule population becomes

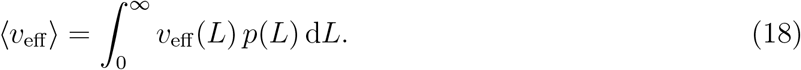

We calculate this effective speed for each motor numerically using the measured distribution *p*(*L*). The end-residence time *τ* was measured for rat kinesin-1 motors to be ≈ 0.5 s [40]. We take this estimate for our K401 motors *(D. melanogaster* kinesin-1) and since, to our knowledge, there is no available data on end-residence times for Ncd and Kif11 motors, we use the same estimate for them (we note that in simulation studies too the end-residence time is typically guessed [9]).

Using this *τ* estimate, the measured distribution *p*(*L*), and the motor speed (*v*) and processivity (λ_*v*_) values from Table 1, we numerically evaluate the relative decrease in the effective speeds of the motors as

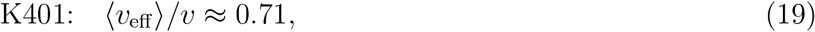

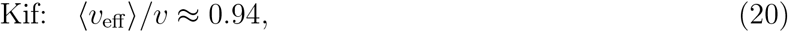

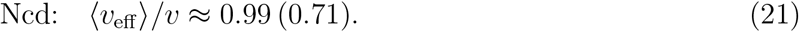

Here we made two estimates for Ncd, first using its single-molecule processivity (≈ 1 step) reported in Table 1, and then the 100-fold increased processivity potentially reached due to collective effects mentioned in the main text. As a consequence of this effective speed reduction, we expect factors of ≈ 1.41, ≈ 1.06 and ≈ 1.01 (1.41) increase in the inferred λ_0_ values of K401, Kif11 and Ncd motors, respectively.

#### 9.15.6 Broader spread of the tubulin profile

In this section, we discuss the feature of a broader tubulin distribution in greater detail. To gain analytical insights, we first consider an idealized scenario where the motor profile can be represented as an exponential decay with a constant offset for the free motor population (Fig. S12a). Such a scenario is approximately met for many of our measured motor profiles. Using our modeling framework, we find that the microtubule distribution corresponding to such a motor profile has the shape of a truncated sigmoid (Fig. S12b, see the second part of this section for the derivation). Indeed, microtubule distributions resembling a sigmoidal shape are observed often in our asters (Fig. S10), two examples of which are shown in Fig. S12c.

One notable implication of this analytical connection between the two profiles is that microtubules necessarily have a broader distribution than motors, once the offset levels at the aster edge are subtracted off. To find whether this is a ubiquitous feature of our asters, we introduce radial distances 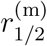 and 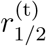 standing for the positions where the motor and tubulin distributions, respectively, are at their mid-concentrations (Fig. S12d). The ratio 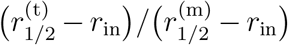, if greater than 1, would then be an indicator of a wider tubulin profile. Calculating this ratio for all of our asters, we find that it is always greater than 1 for all motor types (Fig. S12e), suggesting the generality of the feature.

##### Idealized scenario with an exponentially decaying motor profile

Here, we first derive the analytical form for the tubulin distribution in the idealized scenario where the motor profile can be approximated as an exponential decay. We then demonstrate that, when normalized, this distribution is always broader than the motor distribution.

We start off by writing the motor distribution as

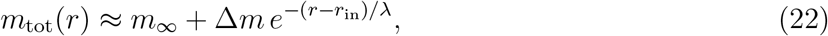

where λ is the decay length scale, *m*_∞_ is the background motor concentration corresponding to the free motor population, and Δ*m* is the amplitude of the exponential decay. Next, using Eq. 6 as well as the definition *m*_tot_(*r*) = *m*_b_(*r*) + *m*_f_(*r*), we set out to obtain the distributions of bound and free motor populations. From Eq. 6, we have

**Figure S12:**
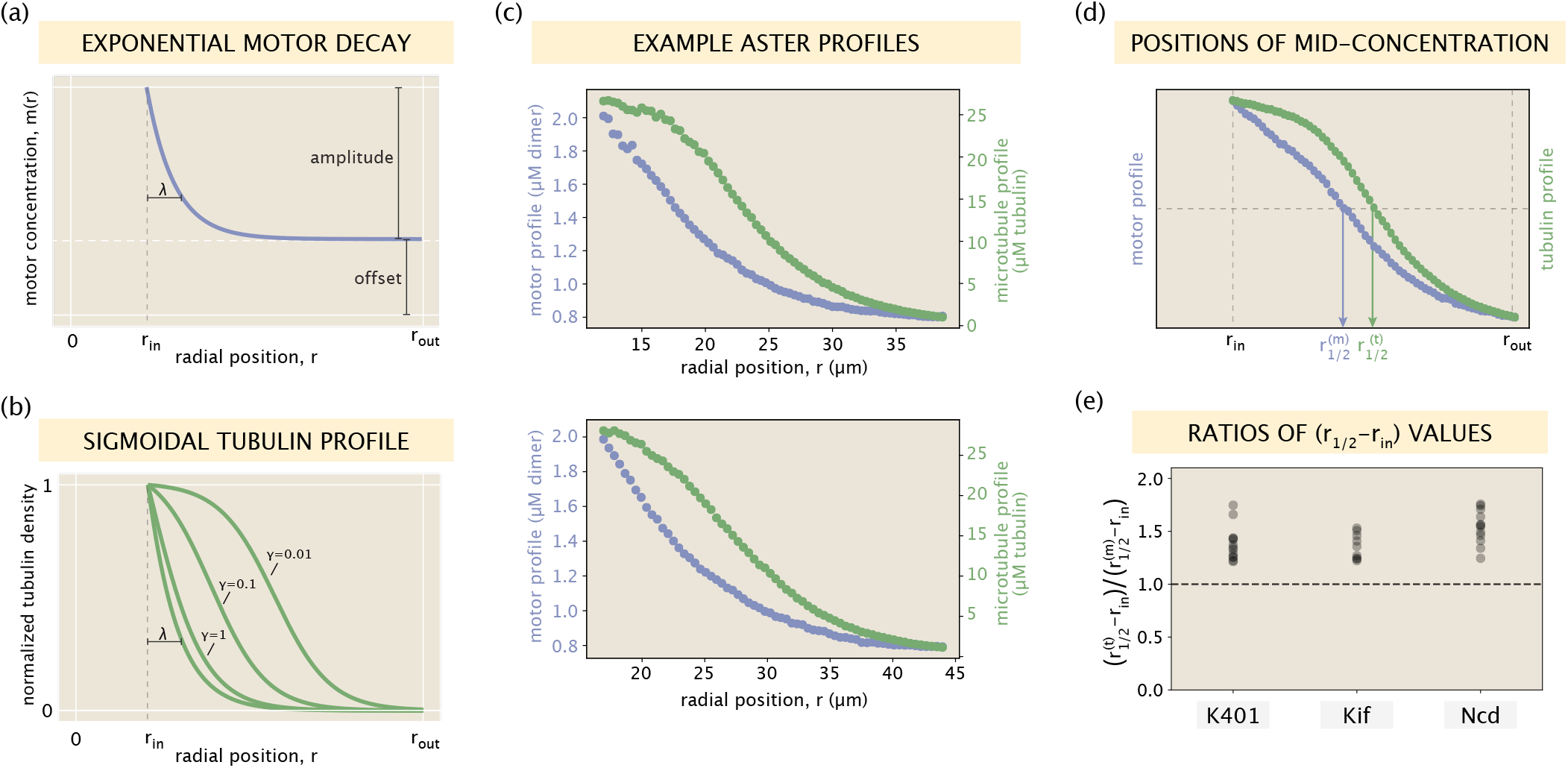
Relationship between motor and microtubule distributions. (a) An idealized exponentially decaying motor profile with a constant offset. (b) Set of sigmoidal tubulin profiles corresponding to the exponentially decaying motor profile. The precise curve depends on the shape parameters of the motor profile and the motor type via an effective constant *γ* (see SI section 9.15.6 for details). (c) Two example profiles from Ncd asters that resemble the setting in panels (a) and (b). Blue and green dots represent measured concentrations of motors and microtubules, respectively. (d) Radial positions in the [*r*_in_,*r*_out_] interval where the motor and tubulin concentrations take their middle values. (e) The ratio 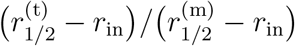 calculated for all of the aster profiles.

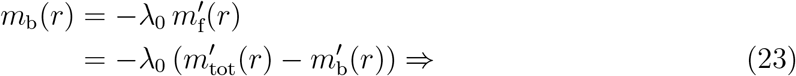

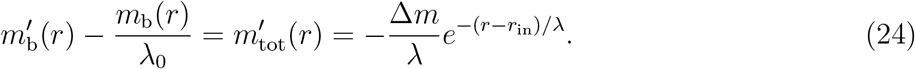

Solving for *m*_b_(*r*), we find

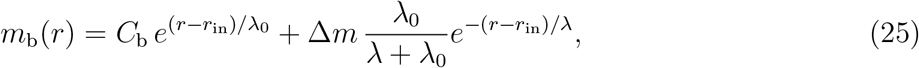

where *C*_b_ is an integration constant. Because the approximation Eq. 22 applies to a finite radial interval *r* ∈ [*r*_in_,*r*_out_], the constant *C*_b_ is generally nonzero. It specifies the relative contributions of free and bound motor populations to the total motor distribution *m*_tot_(*r*).

The free motor population is found by simply subtracting Eq. 25 from Eq. 22, that is,

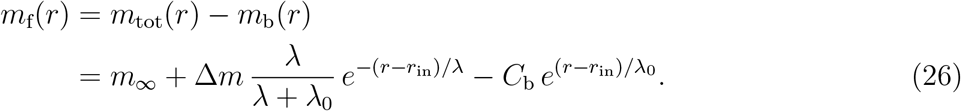

Having obtained expressions for the two motor populations (bound and free), we now recall Eq. 7 that relates these two populations through the local tubulin density. Using Eq. 7, we find the tubulin density as

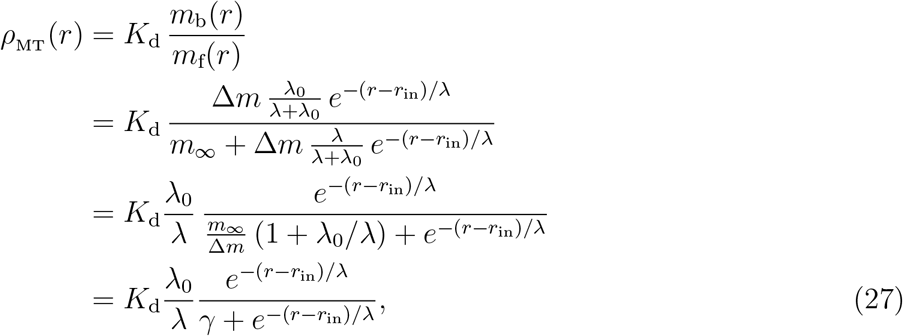

where we introduced the effective parameter *γ* ≡ (*m*_∞_/Δ*m*)(1 + λ_0_/λ). Eq. 27 represents a partial sigmoid, the precise shape of which in the *r* > *r*_in_ region is defined through the parameter *γ* (Fig. S12b).

To formally demonstrate that the tubulin profiles predicted in Eq. 27 are necessarily broader than the motor profile, we first normalized them after subtracting off the concentration values at the outer boundary, namely,

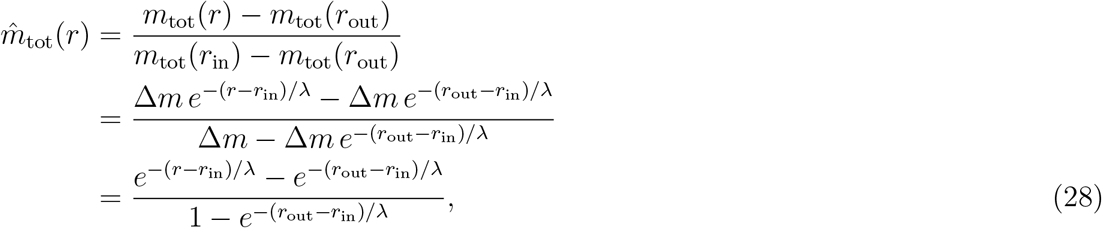

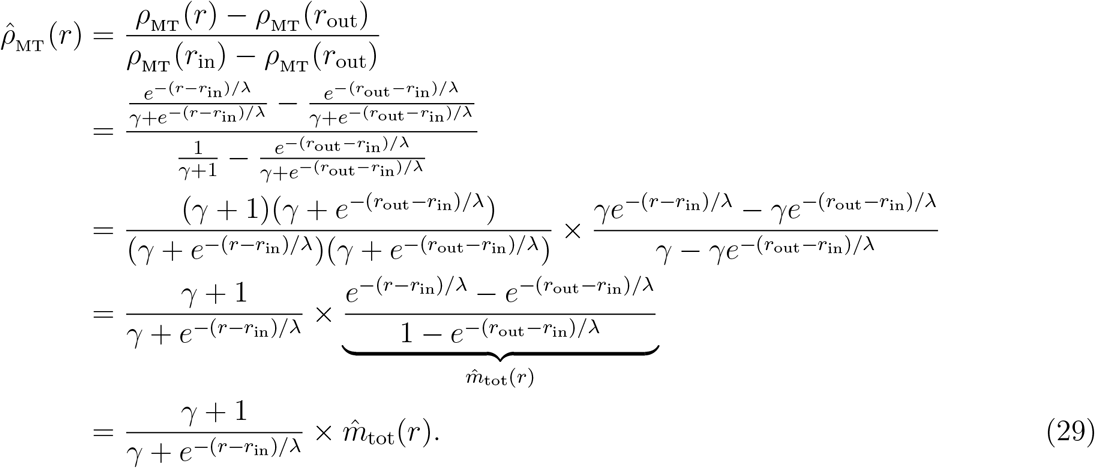

The local ratio of normalized tubulin and motor densities then becomes

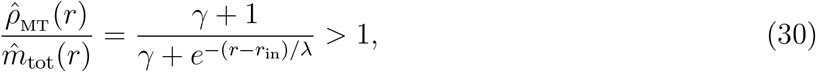

which is always greater than 1 in the *r* > *r*_in_ region. This is indicative of the ‘shoulder’ that the normalized tubulin profile often forms over normalized motor profile and demonstrates the broader spread of the tubulin distribution in this idealized setting.

##### Relative widths of the two distributions

We can see from Fig. S12e that the relative widths of the motor and microtubule distributions differ most for Ncd motors (median ratio ≈ 1.55), while for K401 and Kif, the widths are more comparable (median ratio ≈ 1.35 for both motors). Here we offer an explanation for this difference between the motors using the analytical insights developed earlier in the section.

Specifically, Eq. 30 and Fig. S12b suggest that lower values of *γ* correspond to broader microtubule distributions. Substituting *r* = *r*_in_ in Eq. 27, we can write γ as

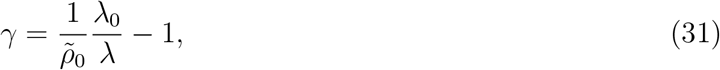

where 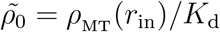 is the microtubule concentration near the core in units of *K*_d_.

From the concentration profiles in Fig. S10, we can estimate the motor decay length scale to be λ ≈ 20,15,10 μm for K401, Kif11, and Ncd, respectively. Then, from our model fitting procedure, we inferred λ_0_ ≈ 40,15, 20 μm, resulting in length scale ratios λ_0_/λ ≈ 2,1, 2 for the three motors. Lastly, again inspecting the profiles in Fig. S10, we find the microtubule concentrations near the core in *K*_d_ units to be 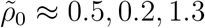. Note that the microtubule concentration near the core (in *K*_d_ units) is the highest for Ncd. In the final step, we substitute these estimated values for λ0/λ and 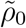 into Eq. 31, and evaluate *γ* for the K401, Kif11, and Ncd, respectively, to be *γ* ≈ {3, 4, 0.3}.

This matches well with the intuition of our simple analytical study (Fig. S12b) and the observed ratios of distribution widths (Fig. S12e). Namely, large values of *γ* for K401 and Kif11 (≈ 3 and 4, respectively) suggest a closer correspondence between the normalized motor and microtubule profiles, while the lower values of *γ* for Ncd (≈ 0.3) suggests a wider microtubule distribution, as was observed in our asters.

## 10 Supplementary Movies

**Movie S1 (separate file)** Activation of K401-Ncd tetramers leads to microtubule bundle formation. Microtubule fluorescence following activation of K401-iLid and Ncd-micro with a 600 μm diameter excitation region is shown.

**Movie S2 (separate file)** Aster formation by K401-K401 and K401-Ncd tetramers. There is approximately a 1:4 ratio of K401-K401 to K401-Ncd tetrameters. Microtubule flourescence is shown.

**Movie S3 (separate file)** Aster formation by K401-K401 and K401-Ncd tetramers. Approximately a 1:2 ratio of K401-K401 to K401-Ncd tetramers is used. Microtubule fluorescence is shown.

**Movie S4 (separate file)** Failed contraction by Ncd-Ncd and K401-Ncd tetramers. There is approximately a 2:1 ratio of Ncd-Ncd to K401-Ncd tetramers. Microtubule flourescence is shown.

**Movie S5 (separate file)** Aster formation by Ncd-Ncd and K401-Ncd tetramers. Approximately a 4:1 ratio of Ncd-Ncd to K401-Ncd tetramers is used. Microtubule fluorescence is shown.

